# Taste matters: Mapping expectancy-based appetitive placebo effects onto the brain

**DOI:** 10.1101/2023.02.14.527858

**Authors:** Iraj Khalid, Belina Rodrigues, Hippolyte Dreyfus, Solène Frileux, Karin Meissner, Philippe Fossati, Todd Anthony Hare, Liane Schmidt

## Abstract

Expectancies, which are higher order prognostic beliefs, can have powerful effects on experiences, behavior and brain. However, it is unknown where, how, and when, in the brain, prognostic beliefs influence appetitive interoceptive experiences and related economic behavior. This study combined a placebo intervention on hunger with computational modelling and functional magnetic resonance imaging of value-based decision-making. The results show that prognostic beliefs about hunger shape hunger experiences, how much participants value food and food-value encoding in the prefrontal cortex. Computational modelling further revealed that these placebo effects were underpinned by how much and when during the decision process taste and health information are integrated into the accumulation of evidence toward a food choice. The drift weights of both sources of information further moderated ventromedial and dorsolateral prefrontal cortex interactions during choice formation. These findings provide novel insights into the neurocognitive mechanisms that translate higher order prognostic beliefs into non-aversive interoceptive sensitivity and shape decision-making.

## INTRODUCTION

A fundamental aspect of human cognition is the ability to extract patterns from noisy sensory information to form prospective beliefs (expectancies) about the world. Statistical frameworks propose that the brain achieves such integration through a computational process that continuously updates prospective beliefs based on prior beliefs and new belief-confirming or disconfirming evidence^1^. This idea has been shared and challenged by many for centuries. An example is Helmholtz’s concept of unconscious inferences^2,3^, which set the foundations for modern enactivist approaches to cognition and computational neuroscience. Importantly, it plausibly accounts for placebo effects. Placebo effects are a famous example of mind-brain-body interactions, wherein the mere suggestion about the benefits of a treatment can shape interoceptive, exteroceptive, and cognitive outcomes^4–10^.

Much previous work has measured aversive outcomes, such as pain combined with functional magnetic resonance imaging (fMRI), to localize placebo hypoalgesia in the brain (for review^11^). However, placebos, which are inactive sham treatments, can also affect appetitive outcome measures. Research on consumption behavior has shown that identical goods are more highly valued and more strongly encoded in the brain’s valuation system when suggested to be more expensive^12,13^. These results are paralleled by findings that verbal suggestion about caloric ingredients or the homeostatic efficiency of a substance can influence the release of satiety signaling hormones^14^, interoceptive hunger experiences^15^, and digestion-related autonomous nervous system responses^16^. Similar to placebo effects in aversive domains, such appetitive placebo effects are mediated by a participant’s beliefs about positive future treatment outcomes, which have also been coined as expectancies or prognostic beliefs^17^.

Despite ample evidence for appetitive placebo effects, there is, however, no direct empirical evidence for when, where, and how, in the brain, an expectancy-based placebo intervention affects the experience of appetitive interoceptive outcomes, such as hunger and associated value-based decision-making. Addressing these questions is important, because it provides an opportunity to understand the effects of higher-order cognitive factors, such as prognostic beliefs, more broadly and how the brain integrates them to make inferences about bodily signals and shape economic behavior that addresses these bodily signals.

To address this question, we built on literature from decision neuroscience that has shown that a person’s goals can affect economic choice behavior. More specifically, generic models of economic choice propose that the decision process involves valuation and action selection (decision) stages^18^. During valuation, various attributes of alternative choices are integrated into stimulus values that approximate hidden preferences that are then compared during the decision stage to select the most preferred alternative (i.e., with the higher stimulus value). The neural mechanisms of both stages have been well studied and involve regions of the brain’s valuation system, such as the ventromedial prefrontal cortex (vmPFC), and cognitive regulation system, such as the dorsolateral prefrontal cortex (dlPFC)^19–27^.

Furthermore, a person’s goals can influence the valuation stage and its related brain responses through cognitive regulation in the form of attentional filtering and value modulation^19,20,28,29^ of relevant information. However, it is unknown whether, when, and where suggestions about interoceptive states can generate such cognitive regulation of valuation and decision stages in the brain.

Given the reported placebo effects on hunger experiences and that economic decision-making addresses hunger, we used a placebo intervention that involved the administration of an identical drink (water), together with the verbal suggestion that the water either increased or decreased hunger. We hypothesized that the placebo intervention would induce prognostic beliefs about the efficiency of the drink’s effect on hunger and through them, affect the experience of hunger, dietary decision-making, and its cognitive regulation. We combined fMRI with a dietary decision-making task and a time-varying drift diffusion model (tDDM) to assess where and to formalize how and when the placebo intervention affected hunger-addressing economic behavior in the brain during decision-making.

In accordance with our hypotheses, we found that the placebo intervention generated hunger expectancies in both groups, which then determined how hungry participants felt at the end of the experiment and moderated the activation of the medial prefrontal cortex at the time of food choice. Consistent with these expectancy-based placebo effects, participants in the increased-hunger suggestion group valued food more highly and displayed stronger vmPFC activation in response to food value than participants in the decreased-hunger suggestion group. Drift diffusion modeling of choice formation showed that participants in the increased-hunger suggestion group considered the tastiness of the food more strongly and rapidly, whereas participants in the decreased-hunger suggestion group considered the healthiness of the food more strongly and rapidly during the decision stage. The drift weights of these two food attributes on the accumulation of evidence toward a food choice then moderated how strongly the vmPFC and dlPFC interacted during choice formation.

## RESULTS

### Expectancy ratings

We first checked whether the placebo intervention was successful in generating prognostic beliefs about hunger. Of note, these were not placebo effects, but rather the participants’ expectancies of how well the drinks would work. Indeed, the suggestions induced, on average, expectancy ratings for both groups that were significantly different from one (i.e., 1 = no expected efficiency) (t(87)_decreased_ = 20.69, p < 0.001 and t(82)_increased_ = 20.09, p < 0.001, one-sample, two tailed t-test). The groups did not differ (mean expected efficiency on hunger: decreased-hunger suggestion group: 5.66 ± 0.23 versus increased-hunger suggestion group: 5.31 ± 0.22; t(169) = 1.13; p = 0.26, two-sample, two-tailed t-test) (Figure 1a).

**Figure 1.**
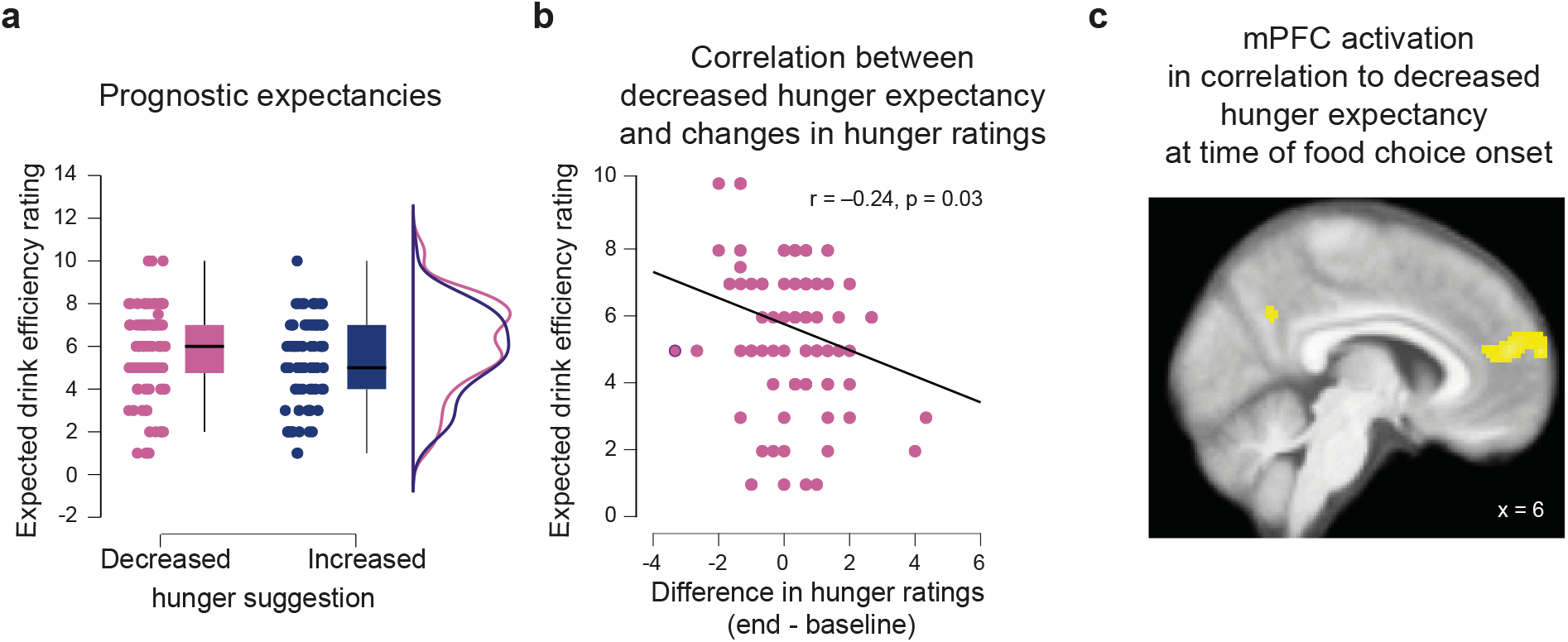
Placebo effects on expectancy ratings. (a) Boxplot graphs displaying the 95% confidence intervals for expectancy ratings for the decreased-(red) and increased-(blue) hunger suggestion groups with the respective distributions of ratings. Dots correspond to individual ratings in each suggestion group. **(b) Correlation between expectancy ratings and the change in hunger ratings from baseline to the end of the experiment**. Each dot corresponds to a participant in the decreased-hunger suggestion group. **(c) Statistical parametric maps of the second-level correlation between brain activation at the time of food choice onset and expectancies about hunger in the decreased-hunger suggestion group**. Voxels in yellow are displayed for visualization purposes at an uncorrected threshold of p < 0.001, and survived p_FWE_< 0.05 family-wise error correction at the cluster level. Activation was taken at the local maxima (MNI x=6) on the sagittal slice, showing the extent of the activation in mPFC from the superior frontal gyrus to the anterior cingulate cortex. SPMs are superimposed on the average anatomical brain image from 57 participants.

Moreover, the more participants in the decreased-hunger suggestion group expected the drink to efficiently decrease their hunger, the less their hunger increased over the course of the experiment (Pearson’s R = -0.24, p = 0.03, Figure 1b). By contrast, this correlation was non-significant among participants in the increased-hunger suggestion group (Pearson’s R = -0.08, p = 0.44).

### Expectancy encoding in the brain at the time of food choices

We then investigated where in the brain these prognostic beliefs were encoded and found that at time of choice, they correlated significantly with activation of the superior frontal part of the medial prefrontal cortex (mPFC) extending into the anterior cingulate cortex (MNI = [2, 58, 22], p_FWE_ < 0.05, cluster corrected, Figure 1c, SI Table 1). This finding was again significant only for the participants assigned to the decreased-hunger suggestion group. The more these participants expected the water to decrease their hunger, the more the mPFC was activated at the time of food choice. No significant moderation of choice-related brain responses was observed for the increased-hunger suggestion group after correction for multiple comparisons at the cluster level. More fine-grained analysis of the brain mediators of hunger experience at the time of food choice formation are reported in the supplement (SI sections 1 and 2, and SI Tables 2 to 5).

### Placebo effects on hunger experiences

Consistent with the effects on prognostic beliefs about hunger, a two-factor (i.e., testing time point by suggestion group) analysis of variance (ANOVA) revealed a significant main effect of testing time point (F(1, 340) = 62.19, p < 0.001) on hunger ratings, which indicated that participants were hungrier at the end of the experiment than at the beginning in both groups (baseline vs end of the experiment: t(87)_decreased_ = -2.53, p = 0.01 and (t(83)_increased_ = -8.99, p < 0.001, paired two-tailed t-test; Figure 2). Importantly, this effect interacted with the suggestion (F(1, 340) = 16.84, p < 0.001). Participants in the increased-hunger suggestion group reported being hungrier at the end of experiment than participants in the decreased-hunger suggestion group (t(170) = -4.14, p < 0.001, two-sample two-tailed t-test; Figure 2).

**Figure 2.**
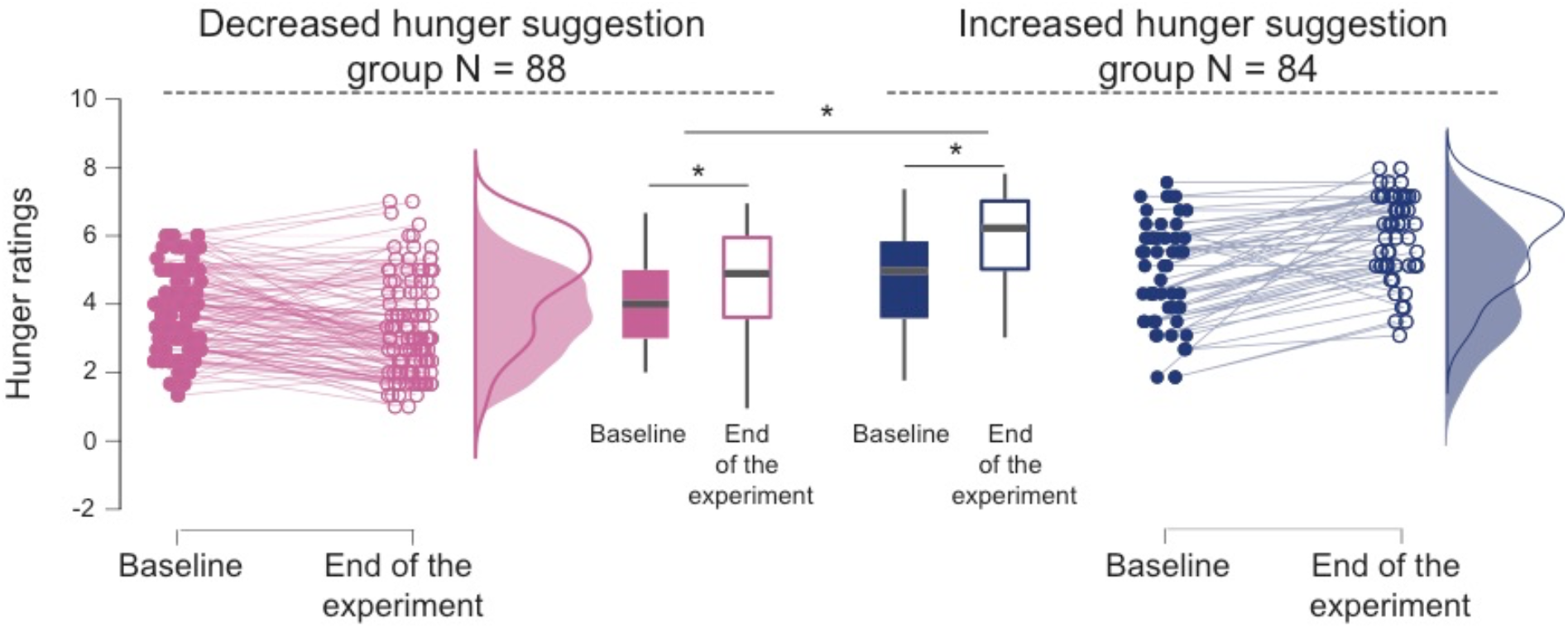
Placebo effect on hunger ratings. Raincloud plots for hunger ratings from baseline to the end of the experiment. Each dot corresponds to a participant’s hunger rating at baseline and at the end of the experiment, with distributions of the hunger ratings displayed on the right side of each plot for the decreased-(red, left-side panel) and increased-(blue, right-side panel) hunger suggestion groups. Boxplots in the middle panel show the 95% CI for both groups at baseline and the end of the experiment. *p < 0.05 for significant within group differences between baseline and the end of the experiment and interaction group (increased > decreased) by the time of the hunger rating (baseline < end of the experiment).

### Placebo effects on food valuation

We then tested the effects of the placebo intervention on hunger-addressing value-based decision-making and related brain responses. Participants in the decreased-hunger suggestion group assigned less value to food stimuli than the participants in the increased-hunger suggestion group (mean SV_decreased_ = 2.09 ± 0.05 versus mean SV_increased_ = 2.28 ± 0.05; t(170) = -2.92, p=0.004, two-sample, two-tailed t-test) (Figure 3a). Consistent with this finding, multilevel general linear regression models of food stimulus values showed a positive prediction by the tastiness of food in both suggestion groups (β_decreased_ = 0.54 ± 0.03, (t(87) = 16.42, p < 0.001, β_increased_ = 0.67 ± 0.03, t(83) = 24.64; p < 0.001, one-sample t-tests, SI Table 6). Importantly, the effect of tastiness on food valuation was much stronger in the increased-hunger suggestion group than decreased-hunger suggestion group (t(170)= -2.81, p = 0.01, two-sample, two-tailed t-test, SI Table 6). These findings are consistent with the more fine-grained analyses of the valuation stage concerning the role and interactions between calorie content, tastiness, and healthiness reported in Supplementary Information Section 3.

**Figure 3.**
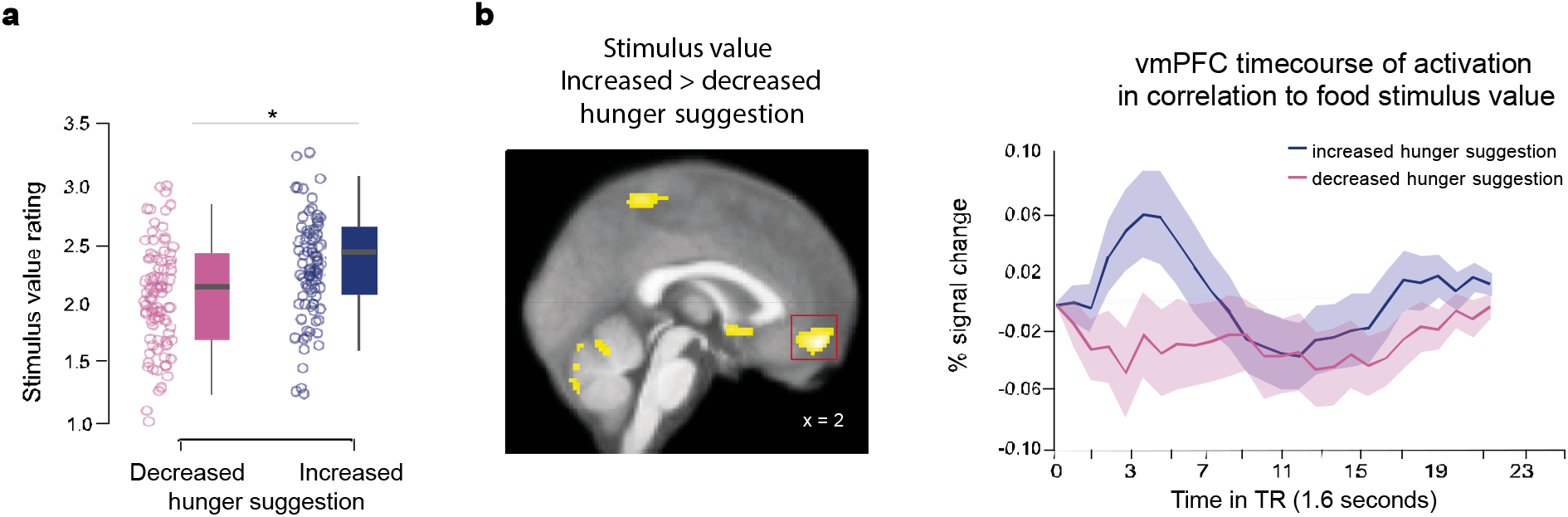
Placebo effect on food preferences and valuation in the brain. (a) Behavioral placebo effect on food valuation. Boxplots showing the 95% CI of the stimulus values assigned to food during the dietary decision-making task for both groups, with jitter elements showing dots for each participant. *p < 0.001. **(b) Neural placebo effect on food valuation**. Statistical parametric maps (SPMs) showing the contrast in brain activation between the increased-versus decreased-hunger suggestion group in response to the stimulus value at the time of the food choice. The significant voxels in yellow survived family-wise error correction based on peak height and cluster (p_FWE_ < 0.05) and are superimposed on the average anatomical brain image. The panel on the right displays peri-stimulus time histograms (psth) extracted from the ventromedial prefrontal cortex global maximum of activation in response to the stimulus value for each suggestion group. The coordinates correspond to the Montreal-Neurological-Institute (MNI) coordinates.

### Placebo effects on valuation-related brain responses

The observed behavioral effects of suggestion on valuation were underpinned by stronger activation of the ventromedial prefrontal cortex (vmPFC), nucleus accumbens, posterior cingulate cortex, bilateral posterior insula, and precuneus in response to stimulus value ratings. Activation of these brain regions correlated more strongly with the food stimulus value in the increased-hunger suggestion group than the decreased-hunger suggestion group (p_FWE_ < 0.05, family-wise error corrected at the cluster level, Figure 3b, SI Table 7). This difference was specific to the encoding of stimulus values. No differences between the two hunger suggestion groups were found for the encoding of tastiness or healthiness in the brain (see SI section 4, SI Tables 8, 9).

### Placebo effects on the decision stage of economic choice

A time-varying drift diffusion model (tDDM) was fitted to the food choices and reaction times and assumed that the decision stage of economic dietary choice is a noisy accumulation of evidence in favor of a “yes” over a “no” food choice. This noisy accumulation of evidence is determined by a series of hidden latent parameters, such as the starting point bias toward a yes or a no choice, sensorimotor integration to select a choice button (i.e., the non-decision time parameter), relative starting time for healthiness (relative to tastiness) to factor in relative to the tastiness of food, respective importance (i.e., drift weights) of healthiness and tastiness, and decision threshold (Figure 4a, SI Table 10). Comparing the two suggestion groups for each of these parameters provides insights into the hidden cognitive processes that are influenced by the suggestion and associated higher order beliefs about interoceptive hunger states.

**Figure 4.**
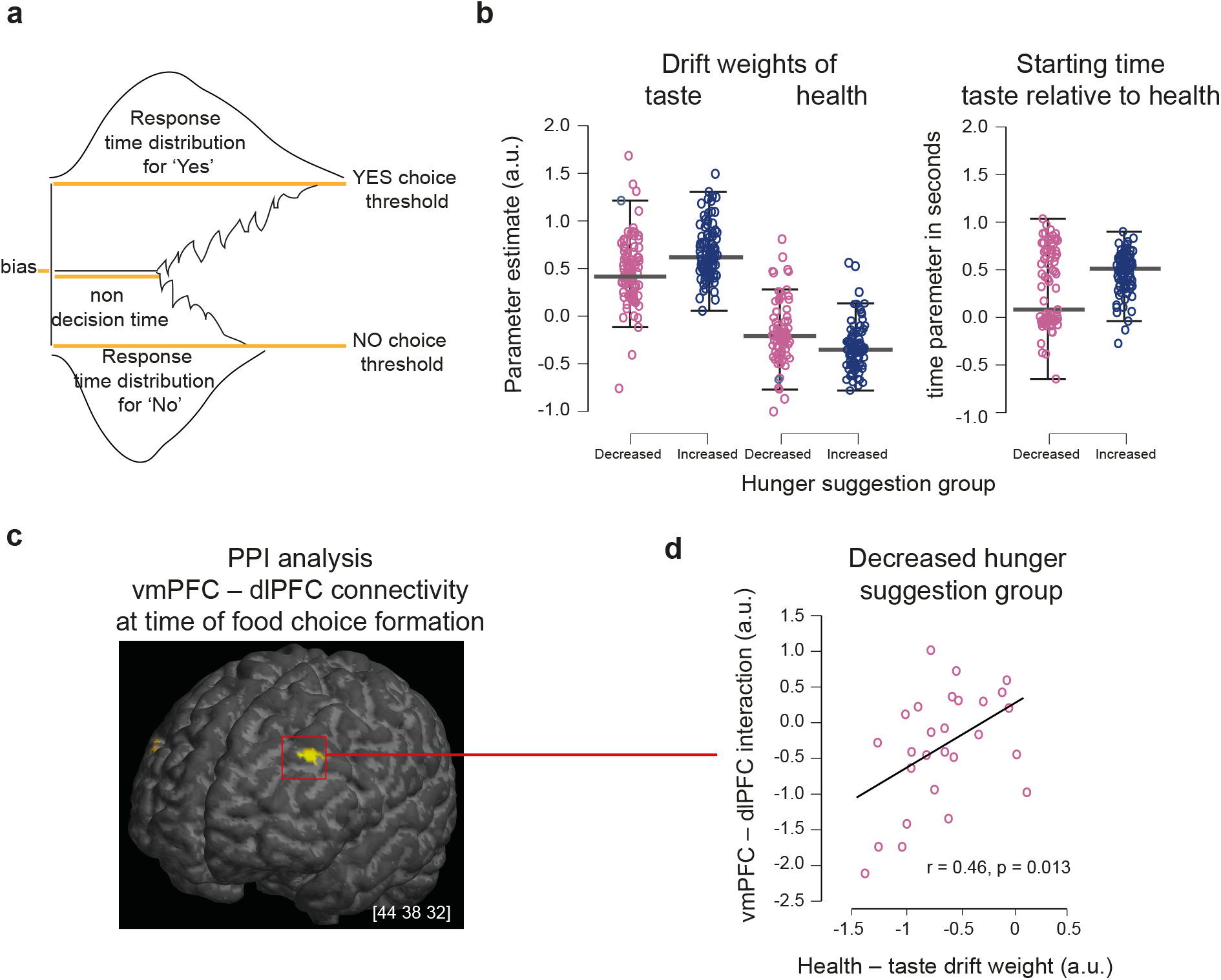
Placebo effects of the integration of tastiness and healthiness in food preferences. (a) Drift diffusion model of food choices. Scheme of the model with free parameters, which involved the starting point bias, non-decision time, choice threshold boundary, drift weights for tastiness and healthiness, and their relative starting time, for which the time is expressed in seconds when healthiness was weighted on the speed of evidence accumulation relative to tastiness. **(b) Individual parameters for tastiness, and healthiness drift weights and relative starting time**. The panels show qualitative differences in the drift weights for tastiness, healthiness, and the relative starting time between the decreased- and increased-hunger suggestion groups. **(c) Psycho-physiological interaction analysis**. SPMs show significant voxels located in the dlPFC that interacted more strongly with the vmPFC seed ROI at the time of making the food choice. The yellow voxels are superimposed on a 3D anatomical brain image and survived small volume correction on the cluster level among the brain regions that were activated in response to interference resolution during the MSIT task. **(d)** The panel on the right shows the correlation between the vmPFC – dlPFC PPI and drift weight of healthiness relative to the drift weight of tastiness obtained in the decreased-hunger suggestion group by fitting food choices and reaction times to a drift diffusion model (DDM).

Choice and reaction time data were fitted in each group using a tDDM and a standard DDM (sDDM; see SI section 5.1 for priors). All chains converged with a Gelman-Rubin convergence diagnostic below 1.5 (psrf = 1). Deviance information criteria were smaller for the tDDM (DIC = 22,873) than the sDDM (DIC = 23,054). Moreover, the log likelihoods of out-of-sample predictions of choices and reaction times observed for even trials by modeled choices and reaction times for odd trials were greater for the tDDM (LL = -742.8) than sDDM (LL = -780.8) (SI section 5.2.). Parameters could be recovered well using an analogous tDDM with a stepwise estimation of the drift rate, as implemented by the deoptim package in R (see SI section 5.3, SI Table 11).

The mean of the posterior distributions of the free parameters of the tDDM revealed that the placebo intervention influenced how much and when participants considered the healthiness and tastiness of the food during the decision process (SI Table 12 and 12a). Note that the health and taste weight coefficients are only defined relative to one another (see equations 2a and 2b). After controlling for tastiness, healthiness negatively influenced the drift rate, but less negatively for participants in the decreased-than increased-hunger suggestion group (mean(PD_decreased_-PD_increased_) = 0.17, PP = 0.95, Figure 4b, SI table 12). On the contrary, tastiness had the opposite effect on the drift rate with a positive weight that was stronger for participants of the increased-than decreased-hunger suggestion group (mean(PD_decreased_ - PD_increased_) = -0.19, PP = 0.99, Figure 4b, SI table 12). The suggestion groups also differed in the relative starting time, which indicated how much earlier (in seconds) the participants considered the tastiness of food relative to its healthiness during evidence accumulation. In both groups the relative starting time was positive, which indicated that tastiness was considered earlier than the healthiness. Importantly, the relative time was shorter for participants given the decreased hunger suggestion than those given the increased hunger suggestion (mean(PD_decreased_-PD_increased_) = -0.24, PP = 0.86, Figure 4b, SI table 12). Similar results were obtained when modeling the choices and reaction times with a tDDM that used a step-wise approximation of the drift rate and was estimated using the deoptim package in R following the procedure of Maier et al. 2020_30_ (SI Table 13, section 5.4).

### Placebo effects on decision stage-related brain responses

To localize where in the brain the placebo intervention affected the decision stage, we searched for the psychophysiological interaction (PPI) of the vmPFC at the time of food choice. We focused on the dlPFC based on previous work providing evidence for the implementation of action selection during the decision process by a vmPFC–dlPFC interaction^19,20,27,31–33^. Consistent with the literature, the PPI analysis indicated significant covariance of the vmPFC with the dlPFC at the time of choice formation for all participants and groups (MNI = [44, 38, 32], p_FWE_ < 0.05 small volume corrected, Figure 4c, SI Table 14 for whole brain activation). Average beta coefficients were extracted from this dlPFC region (MNI = [44, 38, 32]) and correlated with the difference in drift weights between healthiness and tastiness. A significant positive correlation (r = 0.46, p = 0.013) for participants of the decreased-hunger suggestion group (Figure 4d) indicated that the more healthiness scaled the drift rate relative to tastiness of the food, the more strongly the vmPFC – dlPFC interacted during the decision stage of choice formation. The moderation of vmPFC – dlPFC connectivity by healthiness (relative to tastiness) was non-significant for participants in the increased-hunger suggestion group (r = 0.12, p > 0.05).

## DISCUSSION

This study combined computational approaches with brain imaging and behavioral testing to provide insight into the putative mechanisms of placebo effects on appetitive interoceptive hunger experiences and hunger-addressing value-based decision-making. We leveraged a validated placebo intervention that consisted of the administration of an inactive substance (i.e., a glass of water) together with the suggestion that the substance either increases or decreases hunger^15^. In accordance with the results of a previous study^15^, the intervention was successful in generating expectancies about how efficiently the drink would decrease or increase hunger and through them affected how much hunger participants reported over the course of the experiment. Importantly, we provide novel evidence for the underlying putative neurocognitive processes of such appetitive placebo effects. The strength of the prognostic belief in the efficiency of the drink moderated mPFC activation during food choice formation. Consistent with this finding, computational modeling further dissected this effect by showing that the suggestions about hunger influenced the valuation and decision stages of choice formation and the implementation of these two stages of economic choice by the vmPFC and dlPFC.

Past studies have reported mPFC activation in encoding^34^ and computing belief-guided contextual reward expectancies^35,36^ or in representing lower pain expectancies under placebo hypoalgesia^37^. Our results provide novel evidence for recruitment of the mPFC during decision-making as a function of participants’ higher order beliefs about the efficiency of a placebo drink to halt falling energy stores. The moderation was located within the medial frontal gyrus, encompassing Brodmann area 10 and extending into the dorsal anterior cingulate cortex midway between the vmPFC and dlPFC, which have been shown to be part of the brain’s valuation and cognitive regulation system that drives choices under the influence of self-control^19,20,29,38^. Our finding may thus suggest that these participants made more self-controlled food choices. However, dietary self-control is commonly measured by how much participants can forgo a short-term rewarding tasty food in favor of a healthier alternative to meet a goal, such as losing weight or sticking to a healthier diet. In this study, participants were asked to rate their natural food preferences under the suggestion that they had been administered a substance that was designed to influence their hunger. This experimental design is different from that of studies that assess dietary self-control^19,20,28,29,38^ because participants were not instructed to make a choice to meet a specific goal under conflict. Given this difference, it cannot be fully inferred that participants in the decreased-hunger suggestion group were more self-controlled.

However, the placebo intervention may have generated other forms of cognitive and perceptual regulation. For example, perceptual attentional filtering could have played a role during the early phases of the decision-making process and according to the hunger suggestions. Attentional filtering consists of reducing the cost of processing task-irrelevant information, such as overcoming interference during a Stroop task. During dietary decision-making under the influence of higher order prognostic beliefs about hunger states, attentional filtering could consist of considering belief-relevant information more and neglecting belief-irrelevant information. The information under consideration in this example is the tastiness and healthiness of the food. To test this idea, we localized brain activation associated with interference resolution during the multi-stimulus interference task in a supplementary analysis and identified significant activation of the dlPFC. We then found that dlPFC activation at the time of food choice was stronger in the increased-hunger than decreased-hunger suggestion group. After controlling for this contrast effect of hunger suggestion, the dlPFC then positively predicted the increase in hunger relative to baseline and formally mediated the overall suggestion effects on hunger reports. Although this supplementary finding provides only indirect evidence for attentional filtering to be involved when participants make value-based dietary decisions under the influence of their beliefs about hunger, it opens a window toward using other experimental approaches. For example, brain imaging tools such as electroencephalograms could be used, which have more precise temporal resolution to harness the contribution of rapid perceptual regulation mechanisms, such as attentional filtering to placebo effects in interoception.

Another potential cognitive regulation mechanism of the observed placebo effects involves value modulation, which consists of the differential weighting of food attributes during choice formation. We tested this idea directly by building on generic models of economic choices and the finding that the vmPFC is a central hub of the brain’s valuation system that computes expected and experienced values across different decision-making problems and domains^26,39–42^. Our results converge on the finding that suggestions and expectancies can modulate not only subjective self-reports about how hungry participants felt, but also more objective measures of hunger, such as behavioral choices and how the brain encodes values to guide these choices. Computational modelling of choices and reaction times using a time-varying drift diffusion model disentangled several alternative hypotheses about how and when the placebo intervention shaped the decision stage of making the food choice.

Notably, participants in the decreased-hunger suggestion group considered the tastiness relative to healthiness later than the participants in the increased-hunger suggestion group. These computational findings were underpinned at the neural level by moderation of the interaction between the vmPFC and dlPFC during the decision stage. In particular, the more healthiness information weighed on the drift rate, relative to tastiness, the more the vmPFC interacted with the dlPFC during choice formation for participants of the decreased-hunger suggestion group. This is the first study to show that placebo interventions generate higher order beliefs about hunger states and potentially through them, shape how extensively and when in time during the decision stage healthy participants weigh attributes and how the brain encodes this type of cognitive regulation based on value modulation.

Most of the moderation results were specific to participants in the decreased-hunger suggestion group. One may, therefore, reason that this group showed a placebo effect, whereas the increased-hunger suggestion group showed the natural increase in hunger over the course of the experiment. However, this idea needs to be more directly tested in future studies by also randomly assigning participants to a control group that is administered the placebo but without any suggestion about its efficiency in effecting hunger.

Our findings cannot distinguish between several alternative interpretations about the directionality of the generated effects. It may be that the placebo intervention first changed hunger and then the valuation and decision stages of hunger-addressing food choices.Alternatively, the placebo intervention may have generated perceptual and cognitive regulation processes and through them, hunger experiences. Hunger ratings collected in a sub-group of participants before the dietary decision-making task, but after the administration of the placebo, found that the suggestion groups did not yet differ at this time point. Moreover, and as expected, all participants reported being hungrier at the end of the experiment due to longer fasting and explicit weighing of food preferences. The emergence of a group difference in this change in hunger at the end of the experiment, therefore, provides an indication that suggestion-induced cognitive and perceptual regulation may come first and then shape how hungry participants feel. However, more direct evidence is needed to fully disentangle the causal links between these nested effects.

Finally, interoception corresponds to the sensing of bodily states^43^. Here, we used a measure of interoceptive sensitivity by asking participants to self-evaluate their hunger. The question is still open to what extent hunger can be objectively sensed and whether a person’s beliefs about her hunger can affect other dimensions of interoception, such as behavioral and neural measures of interoceptive accuracy and metacognitive confidence in bodily signal detection.

## MATERIALS AND METHODS

### Ethical considerations

The study protocol followed the Declaration of Helsinki and was approved by the local ethics committee. All participants provided written and informed consent and were debriefed at the end of data collection.

### Participants

In total, 188 female participants (mean age = 34.9, sem = 1.02 years, SI Table 15) were recruited for the study via public advertisement in the Paris area.

Participants were screened for right-handedness, normal to corrected-to-normal vision, no history of substance abuse or any neurological or psychiatric disorder, and no medication. Participants of the fMRI experiment were additionally screened for the absence of metallic devices. All participants were tested in the morning between 8 am and 12 pm after overnight fasting. Participants were asked to fast overnight and to not drink tea or coffee at least 2 h before arriving for the experiment. Inclusion was restricted to female participants to minimize gender influences on dietary self-restraint^44^. Participants were paid 15 euros for their participation in the behavioral experiment and 60 euros for their participation in the fMRI experiment.

Exclusion criteria were a baseline hunger rating < 2 (no hunger), pregnancy, claustrophobia, permanent make-up or metallic implants that were not reported at time of recruitment, and technical problems with the fMRI scanner. Based on these exclusion criteria, 16 participants were excluded from the data analysis due to problems with the fMRI scanner (n = 1), not being hungry after overnight fasting at baseline (n = 5 in the decreased-hunger suggestion group and n = 10 in the increased-hunger suggestion group).

After these exclusions, data from 88 participants in the decreased-hunger suggestion group were compared to data from 84 participants in the increased-hunger suggestion group.

### Hunger ratings

Hunger was assessed by three factors: (1) overall experienced hunger (i.e., How hungry do you feel?), (2) homeostatic hunger (i.e., How much food could you eat right now?), and (3) hedonic hunger (i.e., How pleasant would it be to eat, now?). Responses were given on a 7-point Lickert scale from “1” (“not at all”) to “7” (“very much”) and averaged to a common score across the three questions. Hunger ratings were collected at two times during the behavioral study: (1) at baseline, before the placebo intervention and (2) at the end of the experiment, and three times during the fMRI experiment: (1) at baseline, (2) after the placebo intervention but before starting the fMRI session, and (3) at the end of the experiment.

### Randomization

The probability of being assigned to one of the two suggestion arms was fixed to p = 0.5 and constant for the entire duration of the study. Randomization was performed before participants were enrolled using standard permutation algorithms implemented in MATLAB. The algorithm drew 2 integers of 1 and 2. If the integer was ‘1’, the participant was assigned to the suggestion group 1 (decrease hunger suggestion). If it was ‘2’, the participant was assigned to suggestion group 2 (increased hunger suggestion). To ensure an equal number of participants in each suggestion group the permutation was repeated 63 times for the behavioral pilots and 31 times for the fMRI participants.

### Placebo intervention

All participants were administered a glass of mineral water at the beginning of the experiment (®Eau minérale Evian Naturelle). However, the label on the water bottle was specifically designed to provide information about the water’s ingredients to either decrease or increase hunger (Figure 5). In addition, the experimenter explained the labels and all participants read an information booklet about the water’s ingredients and their respective effects on hunger (for further details on the placebo intervention see supplementary information section 6).

**Figure 5.**
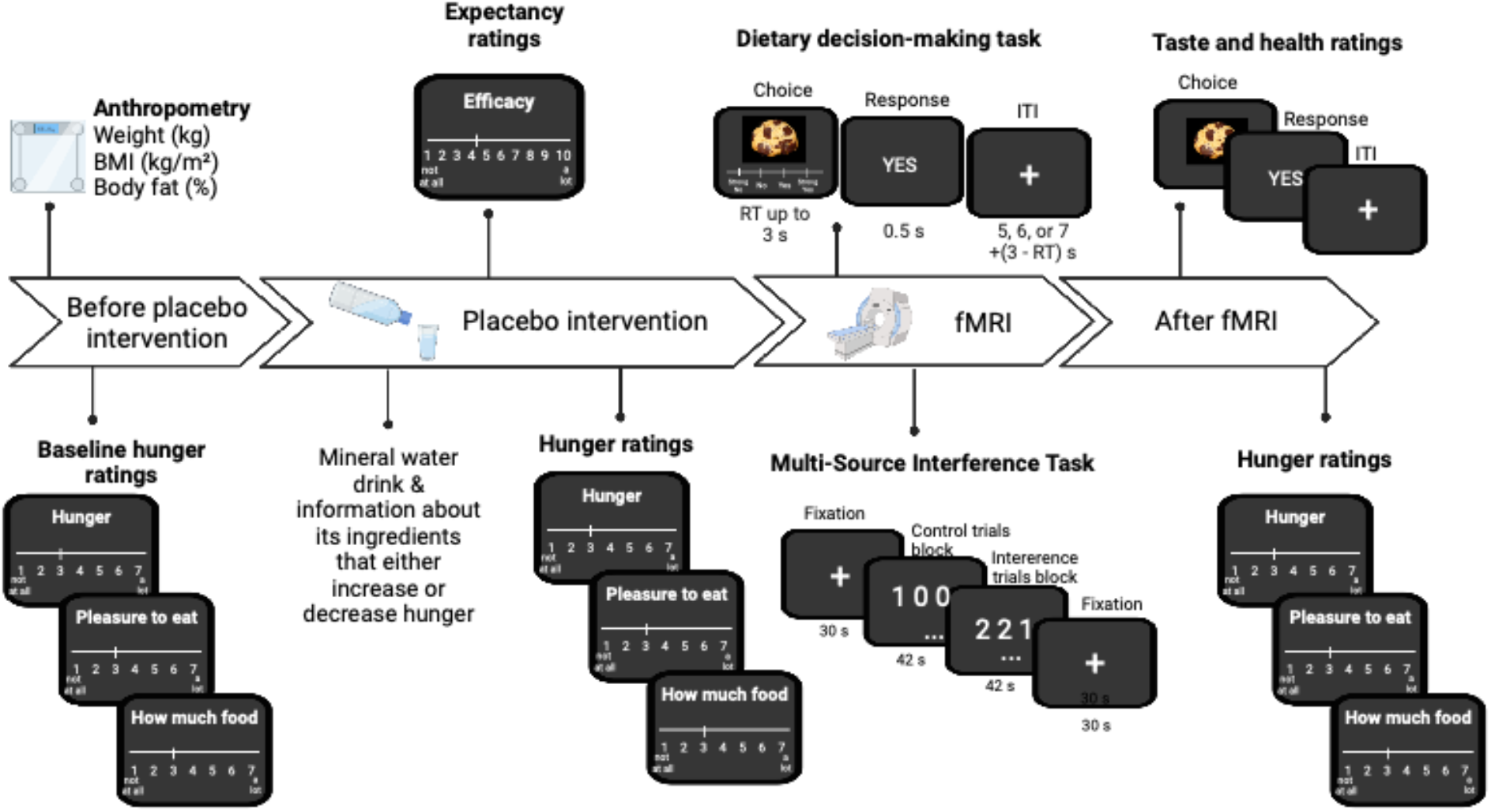
Experimental procedure. The scheme illustrates the temporal organization of the experiment. The black panels correspond to computer screenshots of individual events for the hunger and expectancy ratings given before and after fMRI (outside the MRI scanner), and the dietary decision-making task and multi-stimulus interference task performed during fMRI. For the two fMRI tasks durations are shown in seconds.

Briefly, after rating their baseline hunger, the participants were assigned to the decreased-or increased-hunger suggestion group according to the randomization.

Participants in the increased-hunger suggestion group were told that the drink (water) was enriched with zinc, iron, and plant-based supplements, such as St. John’s Wort, because these ingredients are known for their powerful stimulating effect on appetite through the potentiation of hunger-stimulating hormones, such as ghrelin. By contrast, participants in the decreased-hunger suggestion group were told that the water was enriched with vitamin B12, iron, and riboflavin, because these ingredients had a powerful effect on appetite to curb food cravings through the potentiation of hunger hormones such as leptin.

The experimenter made sure that the participants understood the information about the drink before pouring it into a glass (8.45 oz).

### Expectancy ratings

After drinking the glass of water and before performing the dietary decision-making task, participants rated their expectancy about how efficiently they believed the water drink would decrease or increase their hunger on a 10-point Lickert scale (starting at 1).

### Dietary decision-making task

All participants performed a dietary decision-making task^19,20,29,38^ and a sub-group of 61 participants performed the task during fMRI. The task consisted of the participants choosing whether they wanted to eat snack foods of varying tastiness and healthiness on a trial-by-trial basis. The task counted 200 trials (behavioral pilot) or 152 trials (fMRI sub-group) for a total duration of 20 to 30 min (Figure 5). Each trial started with the display of a food item on a computer screen and participants indicated on a 4-point-Lickert scale, from a strong no to a strong yes, whether they wanted to eat the food item. All food stimuli were selected from a database of 600 food images validated for tastiness and healthiness ratings by 300 participants from a prior Mturk study conducted in-house. The food images were presented on a computer screen in the form of high-resolution images (72 dpi). MATLAB and Psychophysics Toolbox extensions^45^ were used for presentation of the stimulus and recording of the responses. Participants of the fMRI experiment saw the stimuli via a head-coil–based mirror and indicated their responses using a fMRI compatible response box system. The task was incentive compatible, because one food was chosen by chance for consumption at the end of the experiment^19,20^. Participants also rated each food on its tastiness and healthiness using the same 4-point Lickert scale at the end of the experiment.

### MRI data acquisition

T2*-weighted multi-echoplanar images (mEPI) were acquired using a Siemens 3.0 Tesla VERIO MRI scanner with a thirty-two-channel phased array coil. Three echos were acquired for the best compromise between spatial resolution and signal quantity in the orbitofrontal cortex (OFC)^46,47^. To further reduce signal drop out in the OFC, we used an oblique acquisition orientation of 30° above the anterior–posterior commissure line^48^. Each volume comprised 48 axial slices collected in an interleaved manner. To cover the entire brain, the acquisition sequence involved the following parameters: echo times of 14.8 ms, 33.4 ms, and 52 ms; FOV = 192 mm; voxel size = 3 × 3 mm; slice thickness = 3 mm; flip angle = 68°; and TR = 1.25s. Whole-brain high-resolution T1-weighted structural scans (1 × 1 × 1 mm) were acquired for all 61 subjects and co-registered with their mean mEPI images and averaged together to permit anatomical localization of the functional activation at the group level.

### fMRI preprocessing

Image analysis was performed using SPM12 (Welcome Department of Imaging Neuroscience, Institute of Neurology, London, UK). Preprocessing involved the following steps: segmentation of the anatomical image into gray matter, white matter, and cerebrospinal fluid tissue using the SPM12 segmentation tool. The three echo images of each fMRI volume were summed into one EPI volume using the SPM12 Image Calculator^49– 51^. Then, the summed EPIs were spatially realigned and motion corrected, co-registered to the mean image, and normalized to the Montreal Neurological Institute (MNI) space using the same transformation as for the anatomical image. All normalized images were spatially smoothed using a Gaussian kernel with a full-width-at-half-maximum of 8 mm.

### Behavioral analyses

Statistical tests were conducted using the MATLAB Statistical Toolbox (MATLAB 2018b, MathWorks), R (3.3.2 GUI 1.68) within RStudio (RStudio 2022.02.3+492) and JASP (JASP 0.16.4).

### Placebo effects on hunger ratings

For each session (baseline, before MRI, end of experiment) hunger ratings were averaged for the three hunger questions to form one hunger score for each participant. Hunger scores for the baseline and end of the experiment were then analyzed using factorial analysis of variance (ANOVA) with two factors: hunger suggestion group (i.e., decreased hunger coded - 1, increased hunger coded 1) and measurement time (i.e., baseline coded -1, end of experiment coded 1). Post-hoc paired and two-sample t-tests were conducted to characterize the main effects (of group, time) and interaction (group by time). Pearson correlations were conducted for both suggestion groups to assess how the hunger ratings were associated with expectancy ratings.

### Placebo effects on dietary decision-making

Two sample, two-tailed t-tests, along with Bayesian independent sample t-tests, were conducted to compare average stimulus value ratings (SV) between the increased- and decreased-hunger suggestion groups. To further test how the hunger suggestions affected the computation of food preferences at the valuation stage, a multilevel general linear model (GLM) was fitted to stimulus value ratings (SV) following equation (1):

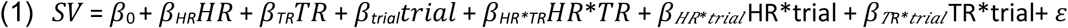

At the individual level, the GLM assumed that food SV was determined by the linear integration of tastiness (TR) and healthiness (HR) attributes of the food, with the rate of integration (beta weights, *β*) varying idiosyncratically between participants. This assumption is consistent with many other decision-making problems and at the core of the valuation phase proposed by models of economic choices. The GLM also included a trial number (trial) regressor to control for fatigue effects and three interactions (TR*HR, TR*trial, HR*trial) to assess how much change occurred in the weights given to the tastiness and healthiness attributes across trials and relative to each other. SV, TR, and HR regressors were mean centered (i.e., coded –2 (strong no), –1 (no), 1 (yes), or 2 (strong yes)). Individual beta weights for each regressor (i.e., *β*) were then fitted into a second level random effects analysis using two-tailed, two-sample t-tests to compare the two suggestion groups. More fine-grained analyses on dietary decision-making are reported in the Supplement (SI table 6).

### Computational modeling

To test how and when suggestions about appetite influenced latent variables of the action selection stage of the decision-making process, SVs were collapsed into binary yes/no choices and fitted together with reaction times by a time-varying drift diffusion model (tDDM). This version of a standard sequential sampling model of action selection has recently been validated by two independent studies for dietary decision-making^30,52^.

The model assumed, similar to traditional sequential sampling frameworks^53–55^, that committing to a choice results from a noisy accumulation of evidence up to a certain threshold in favor of one outcome option (for example, “yes”) over an alternative (for example, “no”). Importantly, the time-varying version of the DDM further assumed that the two sources of evidence, the tastiness (TD) and healthiness (HD) of the food, linearly scaled (ω_tastecp,_ ω_healthcp_) the drift rate (E_cpt_(t)) of evidence accumulation at different times (Time_cp_) within the interval between the reaction time and the non-decision time (DT = RT - nDT). For example, if taste entered the evidence accumulation first, the drift rate (δ_cpt_=E_cpt_(t)) at each timestep (t with dt = 8 ms) was determined by equation 2a_30_:

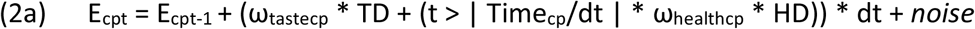

If healthiness entered first, the drift rate at each timestep (dt = 8 ms) was determined by equation 2b:

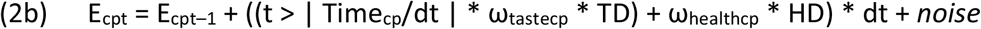

The differences in the tastiness and healthiness ratings for choosing a food item (yes response) versus not (zero response) for a given trial were denoted by TD and HD and respectively scaled the updating of the evidence (the drift rate) by ω_tastecp_ and ω_healthcp_. During the decision time, the time at which the healthiness weighed on the drift rate relative to the time tastiness factored in was expressed by the Time_cp_ parameter. If t > Time_cp_ /dt was false, it equaled 0, whereas if true, it equaled 1. Multiplying one of the two weighted attributes by zero until t > Time_cp_ /dt became true meant that this attribute did not factor in determining the drift rate until a specific time step t. The relative starting time (Time_cp_) parameter was defined by the difference in the time at which healthiness started to scale the drift rate minus the time at which tastiness factored in. A negative Time_cp_ indicated that healthiness influenced the drift rate earlier than tastiness. A positive Time_cp_ indicated that tastiness weighed on the drift rate before healthiness. Note, a Time_cp_ = 0 corresponded to a standard DDM.

Overall, fitting choices and reaction times with a tDDM allowed us to break down the action selection phase into the following hidden latent variables that were then compared between suggestion groups to test how they were influenced by the contextual hunger suggestion: (1) the strength of evidence for a “yes” over a “no” choice (e.g. drift rate), (2) the temporal dynamics of evidence accumulation, assessed by the relative starting time (Time_cp_), (3) how carefully participants made their choices, which was approximated by the decision threshold boundaries, (4) the initial choice bias toward a yes or no food choice, and (5) the non-decision time, which approximated the time taken to initiate a choice and corresponding motor response.

### Model specification

The model was specified using the RWiener package via the run.jags function of the JAGS package in RStudio. More specifically, the tDDM was implemented by a one-dimensional Wiener process, where the state of evidence (dE_t_) at each timestep (dt) evolved stochastically following differential equation (3):

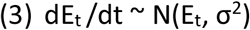

Where E_t_ is the evidence accumulation defined by equations 2a and 2b above. In practice, a stochastic node (y) reflected a certain state of evidence at a specific timestep (dt) (or the predicted choice data and reaction times) and was distributed according to a univariate Wiener distribution:

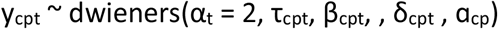

Choice and reaction time (RT) data were coded in a way that “no” food choices were given negative RT values and “yes” food choices positive RT values.

The evidence accumulation started with an initial value of evidence equal to the value of the starting bias parameter (ß_cpt_), which was allowed to vary between participants as a random effect (more details about the priors for ß are provided in SI section 5.1.). The boundary separation parameter (α_t_) was fixed to a maximum value of 2 on a trial-by-trial basis but varied between participants as a random effect. Since each participant was allowed to still have their own boundary separation parameter, the prior for the participant specific alpha (α_cp_) was drawn from a joint normal distribution: α_cp_ = *N*(μ_αcp_, σ^2^_αcp_), with a mean μ_αcp,_ that was itself drawn from a continuous uniform distribution between 0.001 and 2 and a variance σ^2^_αp_ drawn from a gamma distribution with a shape of 1 and a rate of 0.1.

The model estimated the noise in the drift rate (δ_cpt_), which differed on a trial-by-trial basis and between participants. The prior for the drift rate was drawn from each trial (t) from a normal distribution: δ_cpt_ = *N*(E_cpt_, e.p. Ƭ_cp_), with a trial-specific mean that corresponded to the evidence (E_cpt_) accumulated up to this trial following equations 2a and 2b and a variance (e.p. Ƭ_cp_) drawn from a gamma distribution with a shape and rate determined by the error terms of the regression function (see SI section 5.1) that was truncated between 0.001 and 2. The priors for the tastiness (w_tastecpt_) and healthiness drift weights (w_healthcpt_) were defined by uniform distributions between –5 and 5. Both drift weight-free parameters were allowed to vary between participants as random effects.

The non-decision time (τ_cpt_) was also allowed to differ between participants as a random effect, with a mean drawn from a uniform distribution between 0 and 10 and a variance drawn from a gamma distribution with a shape of 1 and a rate of 0.1. Finally, the relative starting time parameter varied between participants as a random effect and was drawn from a joint normal distribution with a mean that itself was drawn from a uniform distribution between -5 and 5 and a variance drawn from a gamma distribution with a shape of 1 and a rate of 0.01.

### Model estimation

Suggestion groups and behavioral and fMRI participant samples were estimated separately. The six free parameters (αp, ß, ω_taste,_ ω_health,_ τ, and RST) were estimated by Gibbs sampling via the Markov Chain Monte-Carlo method (MCMC) in JAGS^56^ to generate posterior inferences for each parameter. We drew a total of 5000 samples from an initial burn-in step and then ran three chains of 10,000 samples. Each chain was derived from three different random number generators with different seeds (see SI table 10). We applied a thinning of 10 to the final sample, which resulted in a final set of 5000 samples for each parameter.

Gelman-Rubin tests were conducted for each parameter to test for the convergence of chains. The potential scale reduction factor (psrf) did not exceed 1.02 for any parameter at the participant or population level, and the deviance (the log posterior) had a prsf ∼ 1.

### Model selection criteria

Choice and reaction time data were fitted using a tDDM and compared to a standard DDM (sDDM) without the relative starting time parameter. Deviance information criteria were used to compare the model fits. The DIC was defined following Gelman et al.^57^ as DIC = 0.5*var/mean(deviance) and was smaller for the tDDM (DIC = 22.873) than sDDM (DIC = 23.054). Parameter recovery, reported in supplementary information, provided estimates that were identifiable (SI table 11). Moreover, posterior checks of predicted and observed choices and reaction time distributions are shown in the supplement (SI figure S9).

### Comparison of free parameters between appetite suggestion groups

To determine whether latent, hidden parameters of the tDDM were different between suggestion groups, the posterior probability of such difference was calculated following equation 4.

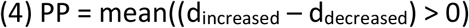

In more detail, a total of 3 posterior parameter distribution chains, counting each 10000 samples, were concatenated for behavioral pilots and fMRI groups for a total posterior distribution over 60000 samples per group-level parameter and suggestion group (e.g., d_increased_, d_decreased_). Then, for each value in each sample a difference was calculated leading to a binary vector of the length 60000. The values in this vector were coded 0 if the difference was smaller then zero (i.e., d_increased_ < d_decreased_) and 1 if the difference was greater then zero (i.e., d_increased_ > d_decreased)_. The mean of those 60000 binary outcomes for each parameter corresponds to the posterior probability that the population parameter distributions (i.e., decreased versus increased suggestion group) differed. Note, except for the healthiness drift weight, all comparisons were made with the prior prediction (H1) that the difference in the posterior parameter distributions between increased and decreased suggestion groups would be greater than zero (SI Table 12 and 12a for group-level mean posterior distributions, and posterior distributions of parameters in each group). In addition, as a sanity check, the individual parameters estimated by a stepwise approximation of the tDDM drift rates and implemented by the deoptim package in R were compared between suggestion groups using Bayesian independent sample t-tests (SI section 5.4 and Table 13).

### Brain imaging analyses

fMRI data were analyzed using Statistical Parametrical Mapping (SPM12, Welcome Department of Imaging Neuroscience, Institute of Neurology, London, UK)^58^. Analogous to behavioral analyses, we searched for suggestion effects on brain responses related to the valuation and action selection phases of dietary decision-making.

fMRI timeseries were fitted using multilevel general linear models (GLM). A first GLM (GLM1) included the following regressors at the first level: an onset regressor at the time of food image display (boxcar duration: reaction time) that was parametrically moderated by the stimulus value, and an onset regressor for missed trials (boxcar duration: 3s). Regressors of non-interest included six realignment parameters (x, y, z, roll, pitch, and yaw) to correct for head movement. Boxcar functions for each trial were convolved with the canonical hemodynamic response function. Individual contrast images for onset choice and the parametric modulator, the stimulus value, were then fitted into a second-level random effects analysis that used two-sample t-tests to localize brain voxels that were activated differently at the time of choice formation and in response to the stimulus value in the decreased-hunger suggestion group (N = 28) relative to the increased-hunger suggestion group (N = 29). Moreover, to test whether brain responses at the time of choice were moderated by expectancies about hunger outcomes, expectancy ratings were added as second-level covariates of the choice onset regressor in a separate GLM (GLM2). GLM2 included the onset regressor at time of choice, with a duration corresponding to the reaction time, and a missed trials onset regressor of a boxcar duration of 3s at the first level. At the second level, one-sample t-tests were used to test how much expected hunger modulated brain responses at the time of choice onset in each suggestion group.

### Time courses

We extracted the activation time courses at the maxima of interest for all reported time course analyses. The response time courses were estimated using a flexible basis set of finite impulse responses separated by one TR of 1.25 seconds.

### Psycho-Physiological-Interaction (PPI) analysis

PPI analysis aimed to localize the brain regions that exhibited choice formation-related functional connectivity within the brain and how such connectivity was linked to free parameters of the tDDM model in each suggestion group. We chose the vmPFC as a seed ROI because it is one of the central hubs of the brain’s valuation system (BVS) that encodes both expected and experienced rewards. Moreover, the vmPFC has been reported to implement action selection in connection with other fronto-parietal brain regions, such as the dorsolateral prefrontal cortex^19,20,27,32^.

Functional timeseries were fitted by a third GLM (GLM3), with three onset regressors at the first level: the time of fixation (duration = 0.5s), choice (duration = reaction time), and missed trials (duration = 3s). Realignment parameters were included as regressors of non-interest to control for head movement. We then extracted average BOLD activity timeseries from a 5-mm sphere centered around the vmPFC ROI (MNI coordinates = [0, 52, -12]) for the contrast choice versus fixation and estimated a fourth GLM (PPI-GLM4), which included a psychological regressor that modeled the choice formation as : reaction time - long boxcars at the time of food choice onset, the physiological regressor of the BOLD activity timeseries of the vmPFC seed region, and the interaction of the psychological and physiological regressors, which was the PPI regressor of interest.Individual betas for this PPI regressor were fitted into a second-level random effects analysis using one sample-t-tests (See whole brain PPI activations in SI table 14).

### Linking tDDM drift weights to vmPFC-dlPFC implementation of evidence accumulation

Beta coefficients from the dlPFC reflected the interaction (in terms of covariance) at the time of choice formation with the vmPFC seed ROI. Beta coefficients from this dlPFC ROI were correlated across participants to the difference between the healthiness and tastiness drift weights from the tDDM (w_healthiness_ – w_tastiness_) using Pearson’s correlations in the both hunger suggestion groups, respectively.

#### dlPFC ROI definition

The PPI with the vmPFC seed at the time of choice formation was small-volume corrected (SVC) using a dlPFC ROI defined by MNI = [40, 42, 26], which was significantly activated during interference resolution measured by the MSIT task (see SI section 1 and SI Table 3). Average beta coefficients reflecting the vmPFC–dlPFC interaction strength at the time of food choice were extracted from a 5-mm radius sphere that was centered around the dlPFC MNI coordinate [44, 38, 32], which survived SVC (p_FWE_ < 0.05, peak height and cluster level).

## Acknowledments

We thank Hilke Plassmann, Bassem Hassan, Mathias Pessiglione, Jean Daunizeau, Nathalie George, Karim N’Diaye and Douglas Lee for helpful comments. This work was supported by core funding from the Paris Brain Institute foundation.

## Data and code availability

Behavioral data, task code, and materials used in the analysis are available on the Open Science Framework website, and can be accessed via the following link: https://osf.io/7j4qs/?view_only=d5c0886514d740c293ebdc14524f37a6

