## Supplemental Information for "Taste matters: Mapping expectancy-based appetitive placebo effects onto the brain"


1. Multi-Source-Interference-Task
   1.1. Behavioral results
   1.2. Whole brain results
2. Brain mediators of placebo effects on hunger experiences
   2.1. Methods: ROI-based single-level mediation analysis
   2.2. Results: ROI-based single-level mediation analysis
3. Additional analyses of dietary decision-making
   3.1. Frequency of response categories during dietary decision-making
   3.2. Responses by tastiness and healthiness categories
   3.3. Regulatory success
   3.4. Relationship between the food stimulus value and calorie content of food
   3.5. Correlation of tastiness to caloric density
4. Additional brain imaging results
   4.1. Correlation of brain activation with the tastiness and healthiness at the time of food choice
5. Computational modeling
   5.1. Priors for the tDDM
   5.2. Out-of-sample predictions of odd to even trials
   5.3. Results when using a stepwise approximation of the drift rate and the tDDM implemented by the Rcpp toolbox in R
   5.4. Parameter recovery
6. Details of the placebo intervention
7. SI Tables 1 – 15

### 1. Multi-Source-Interference task (MSIT)

To localize brain activation related to cognitive regulation, such as attentional filtering of task-relevant information, the participants of the fMRI experiment performed two sessions of the Multi-Source Interference Task (MSIT; Figure 5). The task design followed a previously reported procedure[1–3]. Briefly, the goal of the task was to select the digit 1, 2, or 3 as quickly and accurately as possible by pressing the corresponding response button (i.e., index finger for '1', middle finger for '2', and ring finger for '3'). During congruent control trials, the target number matched its position within zero distractor digits on the computer screen (e.g., 1 0 0 or 0 2 0 or 0 0 3). On the contrary, during incongruent interference trials, the target digit never matched its position within the non-zero distractor digits (e.g., 2 1 2). This condition created a Stroop effect and required allocation of attentional resources to inhibit task-irrelevant information and filter task-relevant information to provide a correct response. Before performing this task within the fMRI scanner, all participants trained for a full session outside the scanner. Behavioral and fMRI analyses of the MSIT task are reported in sections 1.1 and 1.2 respectively.

#### 1.1. Behavioral Results

Four participants were excluded from the behavioral analyses because they misunderstood the task instructions. These four participants reported the place of the target number for the incongruent trials. For example, they pressed the third response button for a "1 1 2" trial, whereas the correct answer was pressing the second button. Two of them performed the training task perfectly but failed during the MRI session despite the instructions being repeated before each session. For the other two participants, performance during the incongruent trials was already close to zero during the training session. For these two participants, it appears that the instructions were not well understood and that the experimenter remained unaware of this during the training session.

For the remaining participants, the effect of interference by task-irrelevant information was measured using two linear-mixed-effect models (LMEs) applied to the two behavioral measurements of interest: performance (i.e., % correct responses) and reaction time (RT, in seconds). In accordance with the literature, participants showed a significantly higher number of errors during the incongruent than congruent trials (SI Table 2), which was indicated by a main effect of trial type (congruent or incongruent) on both performance ($\beta$ = 5.90, t(199) = 5.51, $p < 0.001$) and reaction time ($\beta$ = -0.18, t(199) = -13.01, $p < 0.001$). For both measurements, there was no significant effect of group (performance: $\beta$ = -0.91, t(199) = -0.40, p = 0.69; RT: $\beta$ = 0.002, t(199) = 0.08, p = 0.93) or interaction of hunger suggestion group by type of MSIT trial (performance: $\beta$ = -0.60, t(199) = -0.75, p = 0.46; reaction times: $\beta$ = -0.001, t(199) = -0.23, p = 0.82). Furthermore, we observed a learning effect, as the difference in RT ($\Delta$RT = RT incongruent trials – RT congruent trials) was significantly higher in session 1 than session 2 (t(49) = 3.65, $p < 0.001$, paired-sample two-tailed t-test). Neither performance nor RT differed between hunger suggestion groups (SI Table 2).

#### 1.2. MSIT Brain Imaging Results

A multilevel general linear model contained two onset regressors for the congruent and incongruent trial blocks and six realignment parameters as covariates of non-interest to control for head movement. Individual beta estimates for the contrast between task condition (congruent versus incongruent) were then fitted into a second-level random effects analysis using one-sample t-tests to localize brain responses associated with interference resolution (congruent < incongruent) and the opposite effect (congruent > incongruent).

Whole brain activation for interference resolution (congruent < incongruent) involved the dorsal anterior cingulate cortex (dACC), dorsolateral prefrontal cortex (dlPFC), parietal cortex, and anterior insula, which were more highly activated in the incongruent than congruent trials ($p_{FWE} < 0.05$, family-wise error corrected based on peak height, Figure S1, SI Table 3). In contrast, stronger activation of the brain's reward and valuation system was found in the congruent than incongruent trials. This brain activation involved the vmPFC, posterior cingulate cortex, ventral striatum, hippocampus, and posterior insula (pFWE < 0.05, family-wise error corrected based on peak height, Figure S1, SI Table 4). No significant differences were found for the congruent versus incongruent contrast between the hunger suggestion groups.

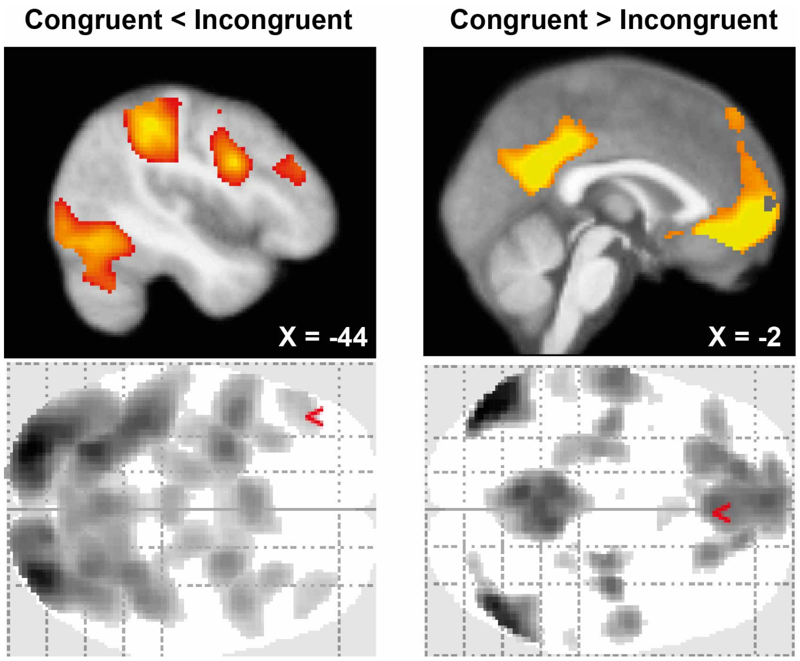


**Figure S1. Brain activation during the MSIT task for interference resolution (congruent < incongruent trials) and control (congruent > incongruent trials).** SPMs are displayed at $p_{FWE} < 0.05$, family-wise error corrected based on peak height. Significant voxels in red and yellow are superimposed on the average anatomical image.

### 2. Brain mediators of placebo effects on hunger experiences

#### 2.1. Methods: ROI-based single-level mediation analysis

To test whether the activation of interference resolution-related brain regions during dietary decision-making mediated the effect of the placebo intervention on hunger experiences, beta estimates were extracted at the time of food choice onset (GLM1) from the following regions of interest: dACC = [-4, 10, 52], right insula = [32, 22, 4], and dlPFC [40, 42, 26] (SI Table 3). The activation in these regions survived small volume correction (SVC) using 10-mm spheres centered on the dACC [3, 10, 47], insula [32, 23, 10], dlPFC [47, 33, 27]), which were reported by Bush et al. 2003[1] to be a robust and reliable neural correlate for interference resolution in humans.

Three ROI-based single-level mediation analyses were conducted, one per ROI, and corrected for multiple comparisons using a Bonferroni corrected p-value of p = 0.05/3 = 0.02 to infer significance. The initial X variable in each mediation model was the suggestion group (coded -1 for the decreased hunger group and 1 for the increased-hunger group). The outcome Y variable was the difference in the hunger ratings (Δhunger = end of experience – baseline), with positive differences indicating greater hunger at the end than at the beginning of the experiment. Path a of the regression for each mediation model tested for a linear effect of suggestion (increased > decreased hunger) on ROI activation at the time of food choice formation. Path b tested for the correlation between this brain activation and hunger by controlling for the effect of suggestion (path a). The path c regression tested for the direct effect of suggestion on hunger and c' for the total effect controlled for the effect of the mediator variable. Finally, mediation was tested by the product of the path a and path b regression coefficients using the formula: $a * b = c - c'$.

Bootstrapping was performed to test the significance of each path coefficient[4,5]. This involved estimating the distribution of individual path coefficients by randomly sampling, by replacement, 10,000 observations from the matrix of [a, b, c, c', a*b] path coefficients. Two-tailed weighted *p*-values were calculated from the bootstrap confidence intervals.

### 2.2. Results: ROI-based single-level mediation analysis

We therefore looked for brain mediators of the placebo effects on the change in hunger ratings from baseline. We reasoned that if the hunger suggestion affects hunger via suggestion-consistent attentional mechanisms, brain regions associated with attentional filtering should mediate such an effect. Attentional filtering can be measured by interference resolution during a Stroop task. We thus first used a localizer Stroop task to identify the brain regions recruited when participants allocated attentional resources to solve interference from task-irrelevant information and to filter task-relevant information (see SI section 1). Average beta estimates for each subject were then extracted from the activation in response to food choice onset in three brain regions of interest that have been reliably reported to become activated during interference resolution[1,3] (SI Table 5). These beta estimates were then used as mediator variables to test the effect of the placebo intervention on hunger ratings. Three single-level ROI-based mediation analyses were conducted, one per brain region (e.g., dACC, insula, dlPFC) that were significantly activated during interference resolution (see SI Table 3 for whole brain activation). The results showed mediation to be significant for voxels located in the dorsolateral prefrontal cortex (dlPFC) ROI (MNI = [40, 42, 26], Bonferroni corrected path a*b, $p = 0.016$, Figure S2). The serial path regressions of this mediation model showed a significant path a effect (beta = 0.49, se = 0.22, p = 0.02), which reflected the univariate effect of the hunger suggestion group (increased > decreased hunger suggestion) on dlPFC activation at the time of food choice. After controlling for the path a effect, dlPFC activation significantly predicted the change in hunger from baseline to the end of the experiment (path b: beta = 0.32, se = 0.11, p < 0.01). Finally, mediation of the suggestion on hunger was significant (path a*b: beta = 0.15, se = 0.09, p < 0.01), with a non-significant total effect of hunger suggestion on the hunger ratings after controlling for the mediator (path c': beta = 0.21, se =0.17, p = 0.26). These findings suggest that the group difference between increased and decreased hunger suggestions on hunger ratings can be formally explained by activation of interference resolution regions located in the dlPFC at the time of making the food choice.

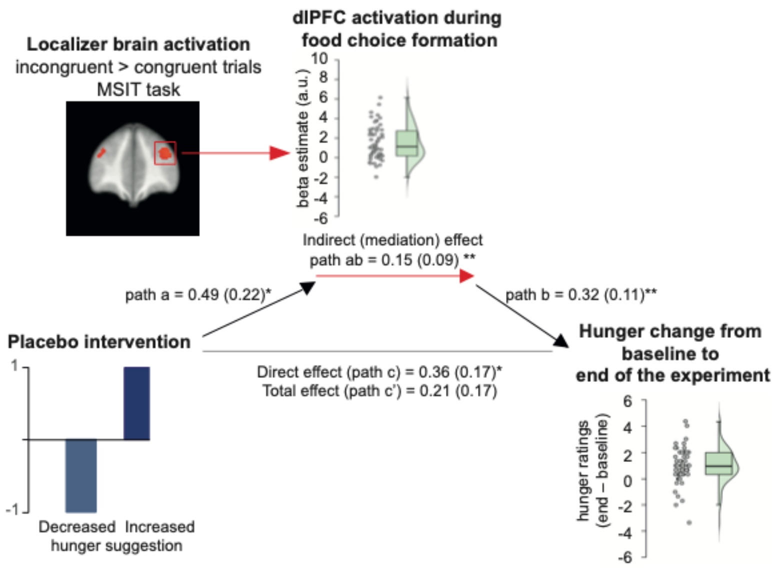


**Figure S2. Brain mediators of placebo effects on hunger.** Single-level mediation path diagram for N = 57 participants for the brain mediator of placebo effects on hunger located in the dlPFC region of interest (ROI) at the time of food choice formation. SPMs in the left panel display the significant voxels of the dlPFC ROI that were activated in response to interference resolution during the Multi-Source Interference Task (MSIT) at $p_{FWE} < 0.05$. The [*x*, *y*, *z*] coordinates correspond to Montreal Neurological Institute (MNI) coordinates and are taken at the maxima of interest. Average path coefficients (a*b (SEM)) between participants denote joint activation in paths a and b at *$p < 0.05$. Note, single-level mediation effects are driven by significant path a and b co-activation of the dlPFC ROI.

### 3. Additional analysis of dietary decision-making

#### 3.1. Frequency of response categories during dietary decision-making

For each food, participants had to rate how much they wanted to eat the food at the end of the experiment using a 4-point Lickert scale with four possible response types: strong no, no, yes, and strong yes. The occurrence of each response type (strong no, no, yes, strong yes) was calculated as the average percentage for a specific type of response:

$$x = \frac{\sum N_{response\ per\ type}}{N_{trial}} * 100$$

The decreased hunger suggestion group had significantly more “strong no” responses than the increased-hunger suggestion group (t(2, 170) = 2.01, p = 0.04, two-sample, two-tailed t-test, Figure S3). On the contrary, the increased-hunger suggestion group responded significantly more often with a “strong yes” than the decreased hunger suggestion group (t (2, 170) = -2.63, p = 0.01, two-sample, two-tailed t-test). No difference was found for the frequency of “no” (t(2, 170) = 0.97, p = 0.33) or “yes” (t(2, 170) = -1.39, p = 0.16) responses between groups.

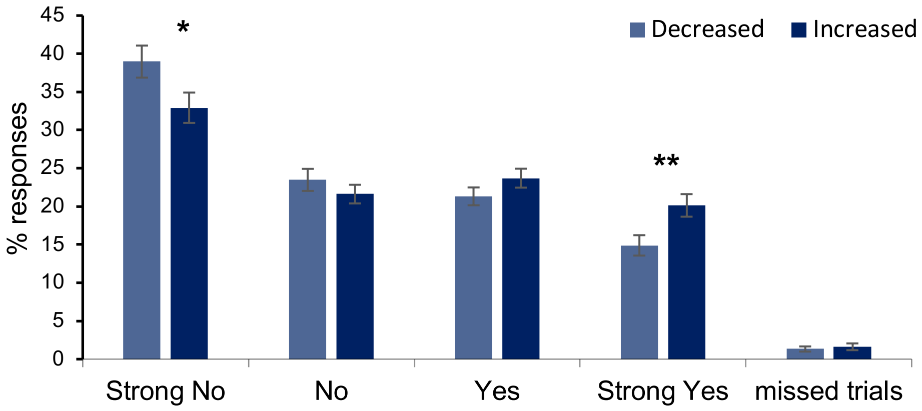


**Figure S3.** Average percentage of each type of response during the dietary decision-making task for the decreased- (light blue bars) and increased- (dark blue bars) hunger suggestion groups. Error bars correspond to standard errors of the mean (SEMs). **P < 0.01, *P < 0.05, two-sample, two-tailed t-tests.

3.2. Responses by tastiness and healthiness categories

Dietary decision-making trials were split into four categories based on the participants’ tastiness and healthiness ratings: Healthy-Tasty, Healthy-Untasty, Unhealthy-Tasty, and Unhealthy-Untasty food choices.

Participants in the increased-hunger suggestion group accepted healthy-tasty food more often ($t(170) = -2.21$, $p = 0.03$, two-sample two-tailed t-test) and rejected unhealthy-tasty food less often ($t(170) = -3.43$, $p < 0.001$, two-sample two-tailed t-test) than participants in the decreased-hunger suggestion group (Figure S4). No significant differences between suggestion groups were observed for healthy-untasty ($t(170) = 1.23$, $p = 0.22$, two-sample two-tailed t-test) or unhealthy-untasty foods ($t(170) = -0.22$, $p = 0.83$, two-sample, two-tailed t-test).

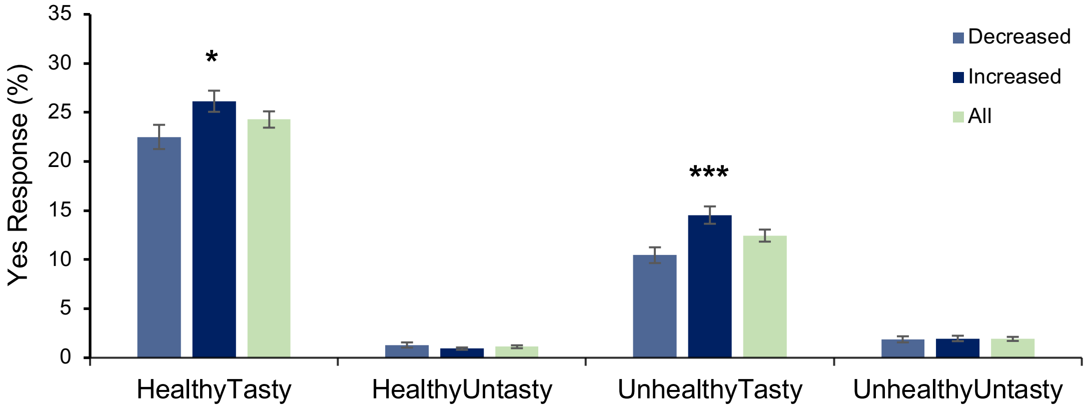


**Figure S4.** Average percentage of binned food acceptance (yes + strong yes responses) as a function of the healthiness and tastiness categories of food in each hunger suggestion group. Error bars correspond to SEMs. ***P < 0.001, *P < 0.05, two-tailed, two-sample t-tests.

#### 3.3. Regulatory success

The regulatory success score (RSS) during dietary decision-making was defined as the sum of accepted healthy-untasty food (yes & strong yes responses) and rejected unhealthy-tasty food (no & strong no responses) over all healthy-untasty and unhealthy-tasty food items using the formula:

$$\text{RSS} = \frac{\Sigma \text{ Yes healthy, untasty} + \text{No unhealthy, tasty}}{\Sigma \text{ healthy, untasty food} + \text{ unhealthy, tasty food}}$$

A significant difference was found between the two hunger suggestion groups ($t(2, 170) = 4.03$, $p < 0.001$, two-sample two-tailed t-test; Figure S5). Participants in the decreased-hunger suggestion group showed a greater RSS (mean $RSS_D = 0.38 \pm 0.02$) than participants in the increased-hunger suggestion group (mean $RSS_I = 0.26 \pm 0.02$).

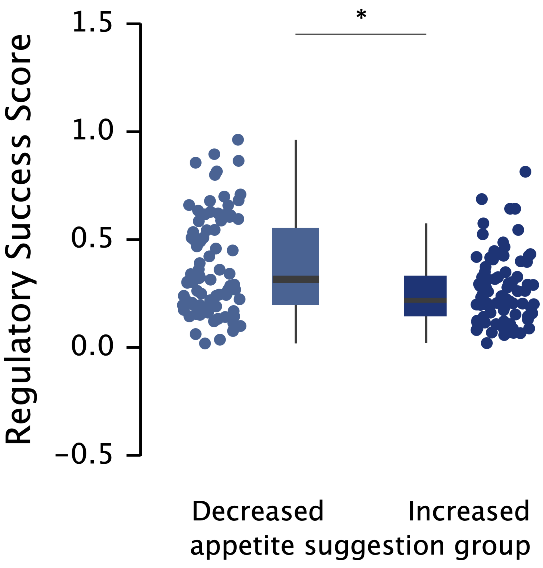


**Figure S5.** Boxplots display the 95% CI for the regulatory success score according to suggestion group (decreased hunger (light blue) or increased hunger (dark blue)), with jitter elements showing dots for the individual score of each participant. Error bars represent the standard error. *P < 0.001, two-sample, two-tailed t-test.

3.4. <u>Relationship between the food stimulus value and calorie content of the food</u>

To determine whether the placebo intervention also affected how calorie content moderated dietary decision-making, we first calculated the calorie density (i.e., calories per gram) for each food item and then tested the correlation between the food stimulus value and calorie density for each participant. We then used one-sample t-tests against zero to test whether the correlations were significant within each group, and found a positive correlation in the increased-hunger suggestion group (average Pearson's R = 0.09, SEM = 0.02, $t(83)_I = 3.89$, $p < 0.001$, one-sample t-test). However, this correlation was non-significant for the decreased-hunger suggestion group (average Pearson's R = 0.04, SEM = 0.03, $t(87)_D = 1.67$, $p = 0.10$, one-sample t-test). The correlation coefficients for the two groups did not significantly differ ($t(2, 170) = -1.29$, $p = 0.20$, two-sample two-tailed t-test).

We then further divided the dietary decision-making trials into two categories, i.e., low-calorie and high-calorie food items using a median split over the calorie densities, for each participant (average median = 1.48, SEM = 0.13, range = 5.99). Overall, we found that the participants did not assign greater stimulus values to high-calorie food than low-calorie food ($t(171) = -1.35$ , $p = 0.18$, paired-sample two-tailed t-test). We obtained the same result when comparing the stimulus values for low-calorie vs high-calorie food for both the decreased- ($t(87)_D$ = -0.39 , p = 0.70, paired-sample two-tailed t-test) and increased-hunger suggestion groups ($t(83)_I$ = -1.60, p = 0.11, paired-sample two-tailed t-test). Importantly, and in accordance with our main findings, participants in the increased-hunger suggestion group assigned more stimulus value to both low- (average $SV_{lowI}$ = 2.23, SEM = 0.04, t(170) = 2.31, p = 0.02, two-sample two-tailed t-test) and high-calorie food ($SV_{highI}$ = 2.30, SEM = 0.05, t(170) = 2.63, p = 0.01, two-sample two-tailed t-test) than participants in the decreased-hunger suggestion group ($SV_{lowD}$ = 2.08, SEM = 0.05, $SV_{highD}$ = 2.10, SEM = 0.05; Figure S6).

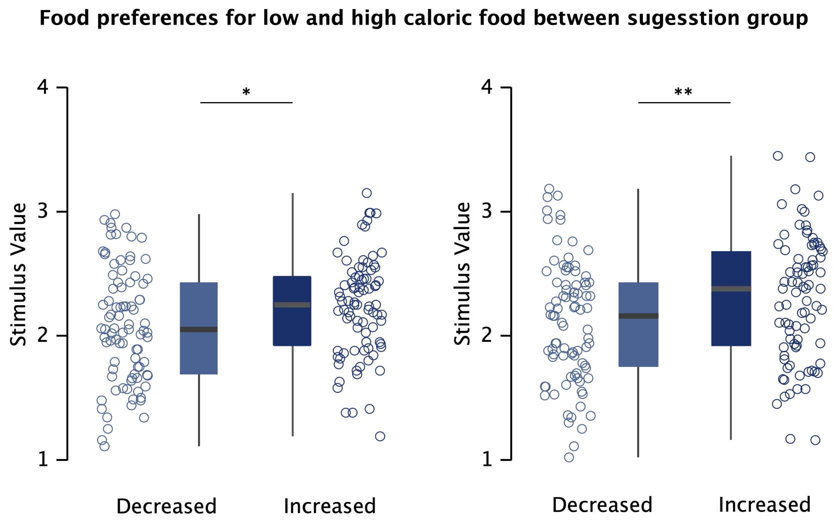


**Figure S6.** Boxplots displaying the 95% CI of stimulus value assigned to low- (left panel) and high- (right panel) calorie food during the dietary decision-making task for the two groups. Individual points correspond to each participant's average SV. Error bars represent the standard error. *P < 0.05, **P < 0.01, two-sample, two-tailed t-tests.

For the entire sample, stimulus values correlated significantly more with high- (average Pearson's R = 0.17) than low- (average Pearson's R = -0.12) calorie food (z = -2.69, p = 0.007, two-tailed Fisher's r-to-z transformation, t(171) = 13.35, p < 0.001, paired-sample two-tailed t-test). This was driven by the increased-hunger suggestion group, which showed a stronger correlation with high- (average Pearson's R = 0.20) than low- (average Pearson's R = -0.13) calorie food (z = -2.19, p = 0.03, two-tailed r-to-z transformation) versus the decreased-hunger suggestion group (average Pearson's $R_{high}$ = 0.16, $R_{low}$ = -0.11, z = -1.62, p = 0.11, two-tailed r-to-z transformation). The differences between groups for the SV correlation with low-calorie food (z = 0.13, p = 0.89, two-tailed r-to-z transformation, t(170) = 0.73, p = 0.47, two-sample two-tailed t-test) and high-calorie food (z = -0.46, p = 0.65, two-tailed r-to-z transformation, t(170) = -1.87, p = 0.06, two-sample two-tailed t-test) were not significant.

#### 3.5. Correlation of tastiness ratings with caloric density

To assess whether the subjective taste ratings correlated with the calorie content of the food items, the calorie density (i.e., calories per gram) was correlated in each participant to the participant's tastiness ratings. Individual Pearson's correlation coefficients were then compared between the increased- and decreased-hunger suggestion groups. Tastiness and calorie density did not correlate in either the decreased- ($t(87)_D = -1.57$, $p = 0.12$) or increased- ($t(83)_I = -0.44$, $p = 0.66$, one-sample t-test) hunger suggestion group. Similarly, there was no difference between the two suggestion groups ($t(2,170)= -0.81$, $p = 0.42$, two-sample two-tailed t-test). This finding indicates that the participants of our sample did not consider high-calorie food to be tastier.

### 4. Additional brain imaging results during dietary decision-making

#### 4.1. Brain activation in correlation with tastiness and healthiness

To localize where in the brain tastiness and healthiness attributes were encoded and whether such encoding differed between the two hunger suggestion groups, we fitted two general linear models (GLMs) to BOLD timeseries with the following regressors: an onset regressor at the time of choice (duration: reaction time) parametrically moderated by tastiness (GLM1) or healthiness (GLM2). Both GLMs also included onset regressors for missed trials (boxcar durations of 3s) and the six realignment parameters as regressors of non-interest to correct for head movement. Individual beta estimates for each regressor were then fitted into a second-level random effects analysis using one-sample t-tests for the entire sample (N=57) and two-sample t-tests to compare the decreased- (N=28) to the increased- (N=29) hunger suggestion groups.

Tastiness at the time of food choice activated the ventromedial prefrontal cortex (vmPFC), posterior cingulate cortex (PCC), precuneus, frontal eye fields, and dorsolateral and dorsomedial prefrontal cortices ($p_{FWE} = 0.05$, family-wise error corrected for the whole brain based on peak height; Figure S7a, SI Table 8). No differences were observed between groups, even with more lenient uncorrected thresholds ($p < 0.001$). Healthiness at the time of food choice was encoded in the bilateral amygdala ($p_{FWE} = 0.05$, family-wise error corrected based on peak height, Figure S7b, SI Table 9). At a lower uncorrected threshold, additional activation was found in the medial prefrontal cortex ($p < 0.001$ uncorrected; Figure S7b, SI Table 9). No differences in brain responses to healthiness at time of food choice were found between hunger suggestion groups. Interestingly, conjunction maps showed the mPFC to correlate with both attributes, but positively with tastiness and negatively with healthiness. This finding suggests that this region integrates the weights of these two attributes differently.

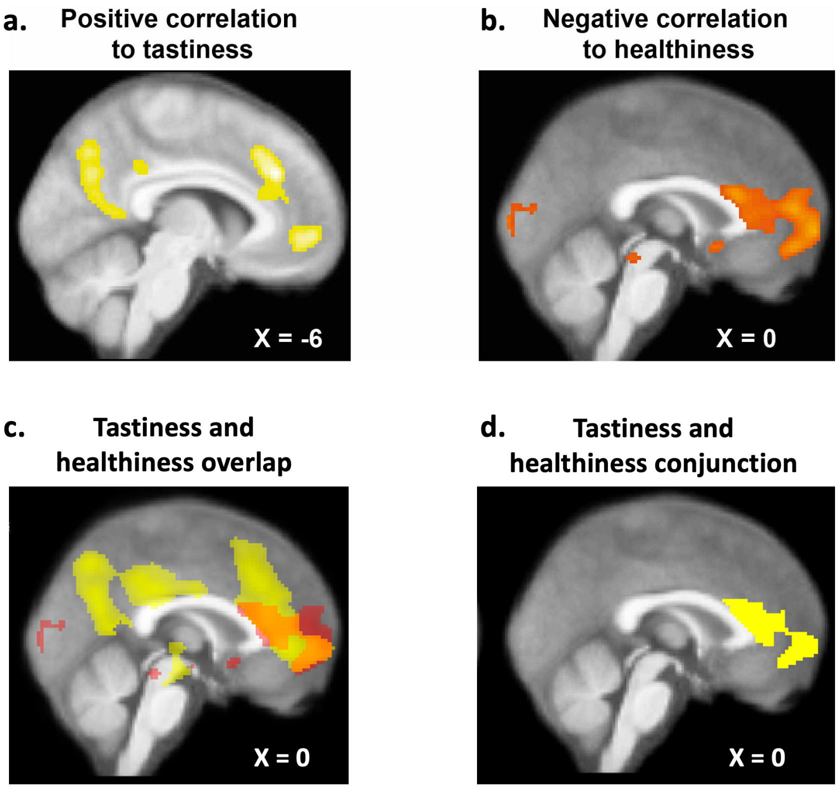


**Figure S7. Brain activation at the time of food choice in response to the tastiness (a) and healthiness (b) of food.** Statistical parametric maps (SPMs) for tastiness are displayed at $p_{FWE} < 0.05$ (family-wise error corrected based on peak height) and $p < 0.001$, uncorrected extent threshold $k = 45$, for healthiness. Voxels displayed in yellow and red are superimposed on the average anatomical image. **(c)** Conjunction map of tastiness (yellow) and healthiness (red) SPMs displayed at $p < 0.001$, uncorrected. **(d)** SPMs for common regions activated in response to tastiness and healthiness displayed at $p < 0.001$, uncorrected.

### 5. Computational modeling

#### 5.1. Priors for each parameter at each level of the hierarchy

We used a hierarchical Bayesian model with three levels of hierarchy: the trial level (t), the participant level (p), and the condition (group) level (C) (Figure S8). The model estimated the most optimal parameters for hidden latent variables of the decision process at each level of the hierarchy, from the lowest trial-by-trial level to the participant and highest condition (group) level. In practice, this involved the model estimating a parameter for the threshold boundary separation ($\alpha_{cp}$), non-decision-time ($\theta_{cp}$), starting bias ($bias_{cp}$), relative starting time ($time_{cp}$), and the drift weights for tastiness ($w_{tastecp}$) and healthiness ($w_{healthcp}$). The model was separately fitted to four groups of participants: the decreased-hunger suggestion fMRI (N=28), decreased-hunger suggestion behavioral pilot (N=60), increased-hunger suggestion fMRI (N= 29), and increased-hunger suggestion behavioral pilot (N = 55) groups.

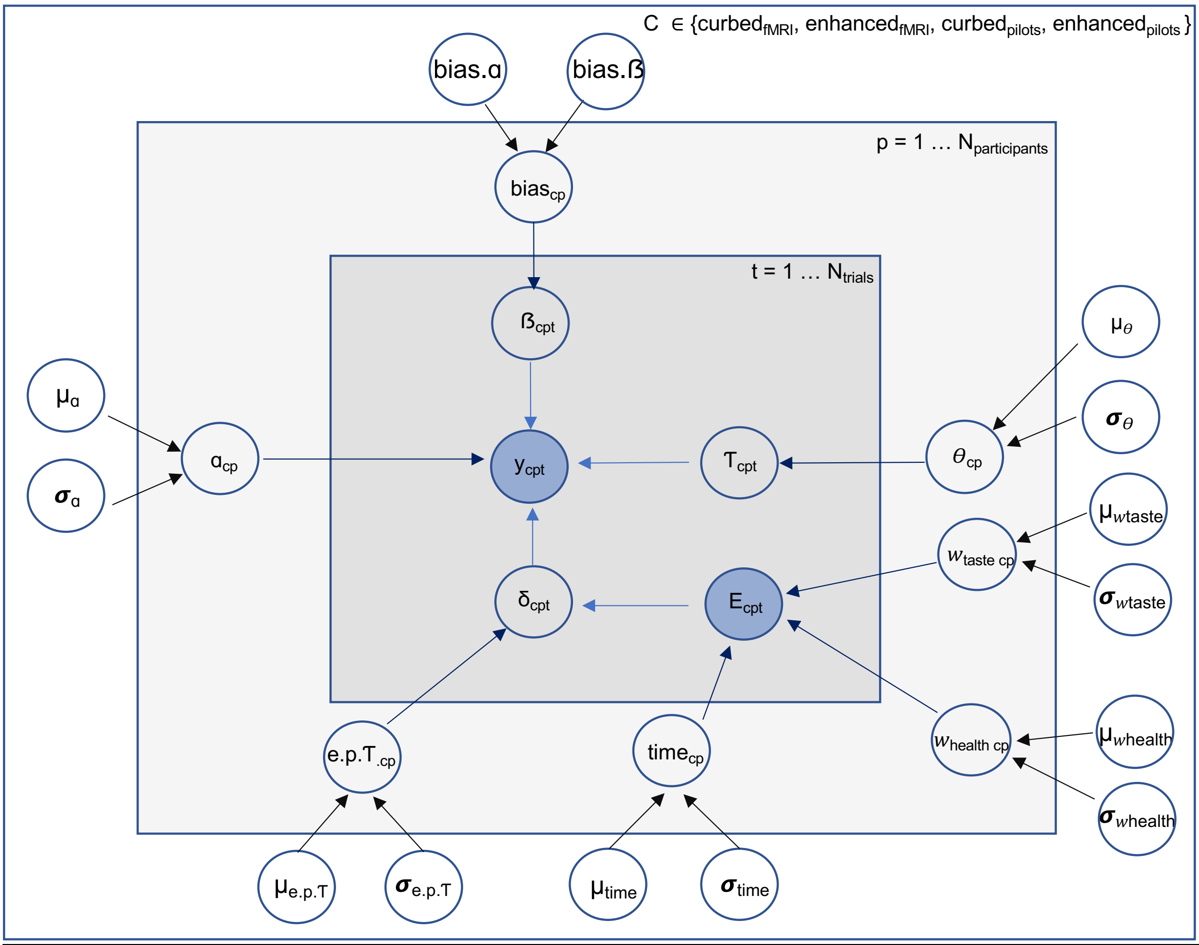


**Figure S8. Scheme of the hierarchical Bayesian tDDM that was fitted to the observed choices and reaction times.** The blue shapes denote the observed variables that varied on a trial-by-trial basis: the evidence, $E_{cpt}$ (defined by equations 2a and 2b, main text), and choice and reaction time data, $y_{cpt}$ (see below). Hidden latent variables are shown as clear shapes at every level of the hierarchy.

#### 5.1.1. Variables on a trial-by-trial level

The priors for predicted choice and reaction time data ($y_{cpt}$) were drawn from each trial from a univariate Wiener distribution using the formula:

$$y_{cpt} \sim dwieners(\alpha_t = 2 \, , T_{cpt}, \beta_{cpt}, \delta_{cpt} \, , \alpha_{cp})$$

Priors for hidden latent variables on the trial-by-trial level were defined by:

- Non-decision time variable $\mathrm{T}_{cpt} = \theta_{cp}$
- Initial starting bias $\beta_{cpt}$ = $\text{bias}_{cp}$
- drift rate $\delta_{cpt}$ ~ dnorm($E_{cpt}$, e.p. $\mathrm{T}_{cp}$)

#### 5.1.2. Variables on the participant level:

The priors for the hidden parameters $\theta_{cp}$, $\text{bias}_{cp}$, and error term e.p. $\mathrm{T}_{cp}$ of the drift rate were defined at the participant level by:

- $\theta_{cp}$ ~ dnorm($\mu_{\theta}$, $\boldsymbol{\sigma}_{\theta}$)
- e.p. $\mathrm{T}_{cp}$ ~ dgamma(e.sG, e.rG)T(0.001, 20)
- $\text{bias}_{cp}$ ~ dbeta(bias. α, bias. β)T(0.01, 0.99)

The priors for the drift weights of tastiness and healthiness were drawn from a univariate normal distribution with the mean μ and precision $\boldsymbol{\sigma}$ using the formula:

- $w_{tastecp}$ ~ dnorm($\mu_{wtaste}$, $\boldsymbol{\sigma}_{wtaste}$)
- $w_{healthcp}$ ~ dnorm($\mu_{whealth}$, $\boldsymbol{\sigma}_{whealth}$)

Priors for the relative starting time ($\text{time}_{cp}$) were drawn from a univariate normal distribution with the mean μ and precision $\boldsymbol{\sigma}$ using the formula:

- $\text{time}_{cp}$ ~ dnorm($\mu_{time}$, $\boldsymbol{\sigma}_{time}$)

The prior for the participant specific threshold boundary separation was defined by:

- $\alpha_{cp}$ ~ dnorm($\mu_{\alpha}$, $\boldsymbol{\sigma}_{\alpha}$)

#### 5.1.3. Variables at the condition (suggestion group) level:

At the highest level of the hierarchy, the condition (group) level, the model assumed flat uniform priors for latent variables, which are also called hyper-condition parameters and corresponded to the mean μ and precision $\boldsymbol{\sigma}$ of the prior distributions for some of the participant level parameters:

- $\mu_{wtaste}$ = unif(-5, 5); $\boldsymbol{\sigma}_{wtaste}$=gamma(1, 0.1)
- $\mu_{whealth}$ = unif(-5, 5); $\boldsymbol{\sigma}_{whealth}$=gamma(1, 0.1)
- $\mu_{time}$ = unif(-5, 5); $\boldsymbol{\sigma}_{time}$=gamma(1, 0.01)
- $\mu_{\alpha}$ = unif(0.001, 2); $\boldsymbol{\sigma}_{\alpha}$=gamma(1, 0.1)
- $\mu_{\theta}$ = unif(0, 10); $\boldsymbol{\sigma}_{\theta}$=gamma(1, 0.1)

- e.SG = pow(e.m.,2) / pow(e.d.,2)
- e.RG = e.m. / pow(E;d.,2)
- e.m. ~ gamma(1, 0.2)T(0.001, 20)
- e.d. ~ gamma(1, 0.5)T(0.001, 20)

- bias. α = $\mu_{bias}$ * bias.kappa
- bias. ß = (1 – $\mu_{bias}$ )* bias.kappa
- $\mu_{bias}$ = beta(2, 2) T(0.01, 0.99)
- bias.kappa ~ gamma(1, 0.5)

#### 5.2. Out-of-sample predictions for odd to even trials

To further check how tDDM performed relative to a standard DDM (sDDM) that did not include the relative starting time parameter, choices and reaction times observed in odd trials were fitted using both models. The obtained parameters were then used to predict the choices and reaction times of the even trials for each participant. We calculated the log likelihoods, which reflected how well modelled choices and reaction times obtained by fitting the tDDM and sDDM to observed data in the odd trials predicted the observed choices and reaction times in the even trials. We found greater log likelihoods for the tDDM for predicting the observed data in the even trials (sum LL = -742.8) than for the sDDM (sum LL = -780.8).

#### 5.3. Results from an alternative estimation method using deoptim in R

We also fitted the observed choices and reaction times with the tDDM using noisy step-wise updating of the evidence accumulation following the procedure of Maier et al. 2020[6]. In more detail, the accumulation started with an initial value of evidence ($E_0$) equal to that of the starting bias parameter (ß) and updated following equations 2a and 2b in discreet time steps of dt = 0.008s until the evidence value $|E_t|$ reached that of the threshold boundary parameter (α). The noise of the step-wise updating was drawn from a Gaussian distribution centered around a mean of zero. The differences in taste and healthiness ratings for a food choice (i.e., a "yes" response) versus a food rejection (i.e., a "no" response) for a given trial were denoted by TD and HD and scaled the updating of the evidence (the drift rate) by $\omega_{taste}$ and $\omega_{health}$, respectively. Once the threshold value was reached, the reaction time RT was computed by $t \times dt + \tau$, with τ corresponding to the sixth free parameter, the non-decision time parameter.

All six free parameters (ß, α, $\omega_{taste,}$ $\omega_{health,}$ τ, and RST (relative starting time)) were estimated with the Rcpp toolbox in R[7] using a differential evolution algorithm[8] and following the estimation procedure reported by Maier et al., 2020[6]. The search space for each of the six free parameters was bounded, with the lower bounds corresponding to (ß = -1, α = 0.6, $\omega_{taste}$ = -2, $\omega_{health}$ = -2, τ = 0.01, RST = -1) and the upper bounds corresponding to (ß = 1, α = 3, $\omega_{taste}$ = 2, $\omega_{health}$ = 2, τ = 1, RST = 1). Tastiness and healthiness ratings were z-scored for all participants within each group. The groups ($increased_{fmri}$, $decreased_{fmri}$, $increased_{pilots}$, $decreased_{pilots}$) were estimated separately.

The results were very similar to those obtained by approximating the drift rate using a one-dimensional Wiener process and estimating the free parameters using Gibbs sampling via the MCMC method in JAGS. Notably, the drift weights for tastiness were greater in the increased- than decreased-hunger suggestion group ($BF_{10}$ = 24.73; SI Table 13) and healthiness factored in earlier in the decreased-hunger suggestion group than in the increased-hunger suggestion group ($BF_{10}$ = 2.20; SI Table 13).

#### 5.4. Parameter recovery

Using a stepwise approximation of the drift rate, we conducted parameter recovery to check whether the tDDM provided parameters that were identifiable and described the data better than any other set of parameters.

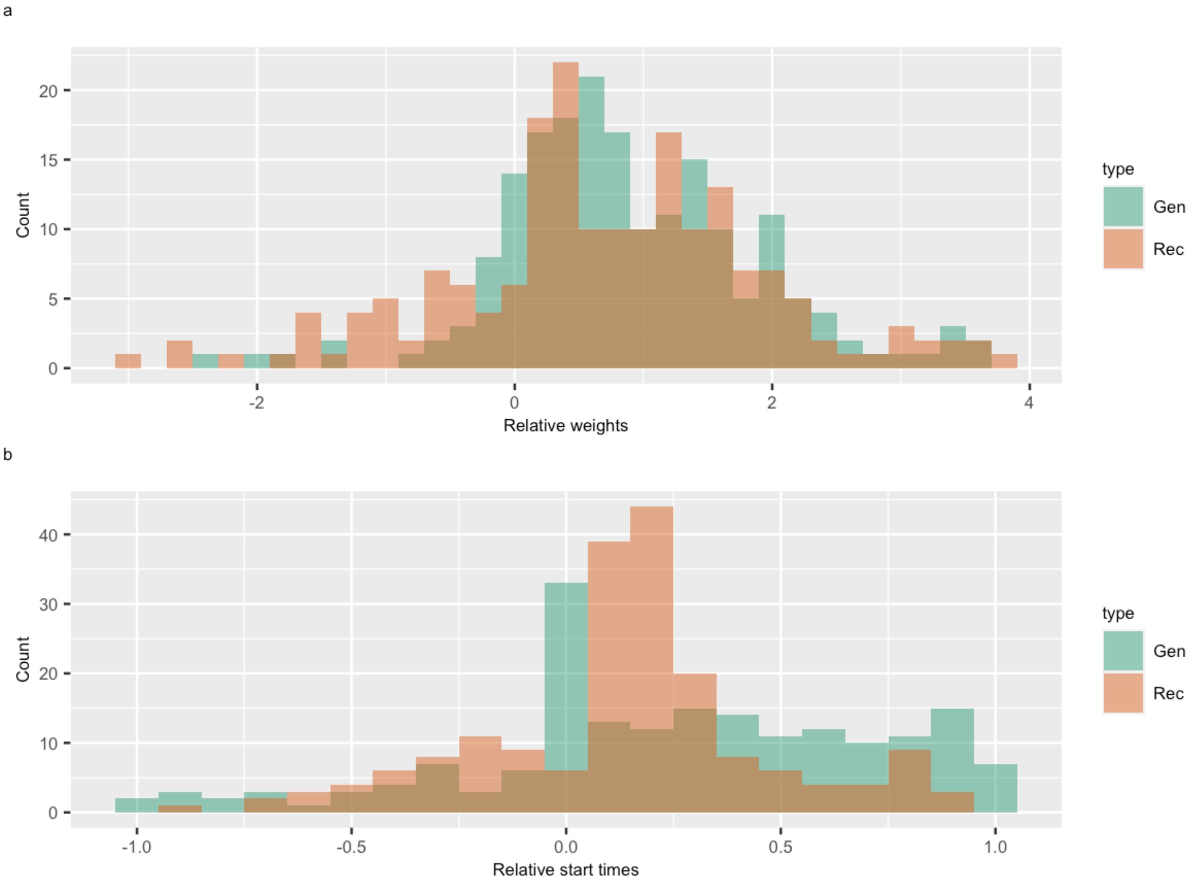


**Figure S9: Histograms for generating and recovering parameters for (a) the relative drift weights and (b) the relative starting times for all 172 participants.**

We first simulated choices and reaction times using the tDDM model, the Rcpp toolbox in R, with the optimal parameter values and observed tastiness and healthiness ratings for the 172 participants. Total trial numbers were determined by each participant's number of observed food choice trials. The simulated data was then fit using the tDDM, which gave a new set of recovered parameters that were then compared to the generating parameters. Parameter recovery was good Figure S8 and SI Table 11). The generating and recovered parameters correlated significantly for the parameters of interest (SI Table 11). Moreover, Bayesian t-tests to search for evidence for a difference between the generated and recovered parameters showed that the evidence for the null hypothesis of zero difference was extremely strong for the two parameters of interest, the relative difference between drift weights ($BF_{01}$ = 0.018), and the relative starting time ($BF_{01}$ = 0.019).

The tDDM gave free parameters that were identifiable in both hunger suggestion groups (Figure S9 and SI Table 11).

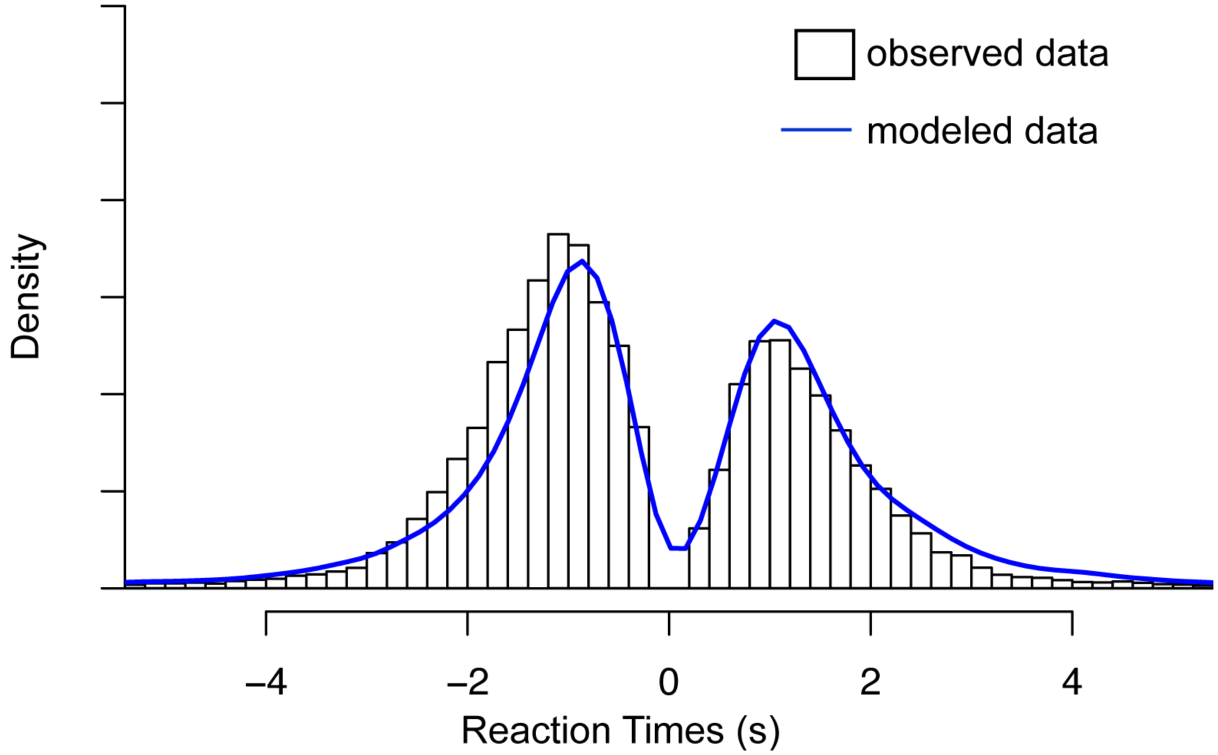


**Figure S10. Modelled and observed choices and reaction times.** Histograms representing the distribution of the observed reaction times across all participants for 'yes' (positive values on the x-axis) and 'no' (negative value son the x-axis) food choices. The blue lines represent model predicted distributions.

#### 6. Details of the placebo intervention

Information about the drink given to the participants assigned to the decreased-hunger suggestion group:

"Before starting the computer/fMRI task, we will ask you to drink a glass of this refreshing, non-carbonated, and neutral-flavored drink, which will help you to focus better on the exercises that you will perform later on the computer/during fMRI. This drink has exceptional nutritional benefits, such as the presence of amino acids, carbohydrates, and traces of elements that are essential for health and brain functioning. Specifically, it has been enriched with riboflavin, also known as vitamin B2, which acts positively on leptin secretion. Leptin is a natural hormone produced by your body that helps to burn fat and reduces food cravings. One glass of this drink is an excellent source of chromium, a formidable anti-sugar element that acts on the metabolism of carbohydrates. It is an insulin cofactor that promotes glucose absorption. Chromium deficiency can cause sugar cravings, with signs of hypoglycemia. Moreover, this water contains vitamins B6, B12, and E, stimulating the development of neural tissue and cell formation, which also act positively on the activity of the neurotransmitter serotonin, known as a well-being hormone involved in mood and concentration. In addition, this water is enriched with mineral salts, such as magnesium and iron, which are essential in case of anemia and a valuable ally for women's health because they provide oxygen to the cells and reduce the feeling of fatigue. This drink is an excellent source of extra food during sports and intellectual endeavors. A 25cl glass covers all the nutritional needs of the next two hours. This drink is natural, gluten-free, and packaged in a protected environment. You will have two minutes to drink a glass of the water. Thereafter, you should wait a few minutes to allow time for the active substance to deliver their effect. We will then ask you to perform a food-choice task."

Information about the drink provided to participants in the increased-hunger suggestion group:

"Before starting the experiment, we will ask you to drink a glass of this refreshing, non-carbonated, and neutral-flavored drink, which will help you to focus better on the exercises that you will perform later on the computer/during fMRI. This drink has exceptional nutritional benefits, such as the presence of amino acids, carbohydrates, and traces of elements that are essential for health and brain functioning. Specifically, it has been enriched with zinc, which promotes digestion, fights against heartburn, and stimulates the taste buds. It also brings together the benefits of medicinal plants such as St. John's wort, known for its stimulating effect on hunger. Its action is based on its ability to send a hunger message to the brain. Several studies have shown that this medicinal plant acts positively on the activity of the neurotransmitter serotonin, known as a well-being hormone involved in mood and concentration. In addition, this water is also enriched with mineral salts, such as magnesium and iron, essential in cases of anemia and a valuable ally for women's health because they provide oxygen to the cells and reduce the feeling of fatigue. This drink is natural, gluten-free, and packaged in a protected environment. You will have two minutes to drink a glass of the water. Thereafter, you should wait a few minutes to allow time for the active substance to deliver their effect. We will then ask you to perform a food-choice task."

### 7. Supplementary tables

SI Table 1. Positive moderation of brain activation by expectancies at the time of food choice in the decreased-hunger group

| Region | Brodmann Area | Side | Size | x | y | z | Peak z-score |
|---|---|---|---|---|---|---|---|
| IpS | 39 | L | 63 | -28 | -80 | 38 | 4.28 |
| mPFC* | 10 | R | 355 | 2 | 58 | 22 | 3.98 |
| AG | 39 | R | 141 | 54 | -66 | 24 | 3.71 |
| dPCC | 31 | R | 53 | 2 | -56 | 34 | 3.48 |

IpS – intraparietal sulcus; AG – angular gyrus; mPFC – medial prefrontal cortex; dPCC – dorsal posterior cingulate cortex. Values are reported at $p < 0.001$, uncorrected, using an extend threshold of $k = 50$. *Regions that survived family-wise error correction based on the cluster level.

SI Table 2. Average performance and reaction times during the MSIT

| | Session 1 | | Session 2 | |
|---|---|---|---|---|
| | **Decreased** | **Increased** | **Decreased** | **Increased** |
| Performance for congruent | 97.61 ± 1.45 | 98.57 ± 0.48 | 92.17 ± 4.49 | 97.92 ± 0.68 |
| Performance for incongruent | 86.11 ± 3.23 | 90.70 ± 1.90 | 82.75 ± 5.04 | 90.63 ± 1.84 |
| $\Delta_{Performance}$ | 11.50 ± 3.00 | 7.87 ± 1.70 | 9.42 ± 3.16 | 7.29 ±1.87 |
| RT for congruent | 0.57 ± 0.03 | 0.54 ± 0.02 | 0.58 ± 0.03 | 0.56 ± 0.02 |
| RT for incongruent | 0.89 ± 0.03 | 0.88 ± 0.03 | 0.89 ± 0.03 | 0.85 ± 0.02 |
| $\Delta_{RT}$ | 0.33 ± 0.02 | 0.34 ± 0.02 | 0.31 ± 0.02 | 0.30 ± 0.02 |

$\Delta_{Performance}$ – difference in performance between congruent trials and incongruent trials ; RT – reaction time ; $\Delta_{RT}$ – difference in reaction times between congruent trials and incongruent trials

SI Table 3. Whole brain activation for the contrast between incongruent > congruent MSIT conditions

| Region | Brodmann Area | Side | Size | x | y | z | Peak z-score |
|---|---|---|---|---|---|---|---|
| V1 | 18 | L | 25142 | -26 | -90 | -4 | Inf |
| SMA | 6 | L | 5164 | -26 | 0 | 50 | Inf |
| mPFC/IFG | 9 | L | | -40 | 4 | 30 | Inf |
| **dACC*** | **32** | **L** | | **-4** | **10** | **52** | **7.49** |
| IFG | 6/9 | R | 782 | 44 | 6 | 28 | 7.04 |
| Thalamus | | L | 965 | -16 | -16 | 14 | 6.71 |
| **ant Insula*** | **13** | **R** | **354** | **32** | **22** | **4** | **6.49** |
| | 13 | L | 232 | -30 | 20 | 6 | 6.61 |
| **dlPFC/MFG*** | **46** | **R** | **303** | **40** | **42** | **26** | **5.50** |
| | 46 | L | 288 | -44 | 38 | 26 | 6.11 |
| Cerebellum | | L | 70 | -24 | -66 | -48 | 6.03 |

V1 – primary visual cortex; SMA – supplementary motor area; IFG – inferior frontal gyrus; mPFC – medial prefrontal cortex; IFG – inferior frontal gyrus; dACC – dorsal anterior cingulate cortex; ant Insula – anterior insula; dlPFC – dorsolateral prefrontal cortex; MFG – medial frontal gyrus; Inf – infinite. Values were taken at pFWE < 0.05 family-wise error corrected based on peak height and cluster. *Regions of interest for the mediation analyses, survived small volume correction using main ROIs reported for the MSIT task by Bush et al. 2003 (dACC [3 10 47], Insula [32 23 10], dlPFC [47 33 27]).

SI Table 4. Whole brain activation for the contrast between congruent > incongruent MSIT conditions.

| Region | Brodmann Area | Side | Size | x | y | z | Peak z-score |
|---|---|---|---|---|---|---|---|
| PPC | 39 | L | 1243 | -46 | -74 | 32 | Inf |
| | 39 | R | 976 | 46 | -70 | 36 | 7.70 |
| PCC | 23/31 | L | 2637 | -6 | -56 | 14 | 7.14 |
| vmPFC | 10 | L | 5474 | -2 | 58 | -2 | 6.89 |
| OFC/vlPFC | 47 | L | 213 | -38 | 34 | -12 | 5.80 |
| NAcc | | R | 30 | 8 | 8 | -12 | 4.85 |
| post Insula | 13 | R | 200 | 36 | -14 | 18 | 6.71 |
| STG | 21/22 | L | 513 | -60 | -14 | -14 | 6.63 |
| | | R | 323 | 54 | -6 | -20 | 6.19 |
| Hippocampus | | R | 130 | 26 | -18 | -20 | 6.30 |
| | | L | 307 | -26 | -20 | -18 | 5.97 |
| Fusiform | 37 | R | 44 | 26 | -36 | -14 | 5.06 |
| MTG | 21 | L | 108 | -56 | -40 | -4 | 5.00 |
| Cerebellum | | R | 64 | 40 | -74 | -42 | 5.32 |
| S1 | 1 | R | 43 | 52 | -8 | 10 | 5.06 |

PPC – posterior parietal cortex; PCC – posterior cingulate cortex; vmPFC – ventromedial prefrontal cortex; OFC – orbitofrontal cortex; vlPFC – ventrolateral prefrontal cortex; NAcc – nucleus accumbens; post Insula – posterior insula; STG – superior temporal gyrus; MTG – middle temporal gyrus; S1 – primary somatosensory cortex, Inf –infinite. All values survived pFWE < 0.05, family-wise error corrected based on peak height.

SI Table 5. Average beta estimates extracted at the time of food choice from regions activated during interference resolution trials of the MSIT

| | **Decreased** | **Increased** |
|---|---|---|
| dACC [-4, 10, 52] | 3.55 ± 0.34 | 3.71 ± 0.22 |
| Insula [32, 22, 4] | 1.95 ± 0.20 | 2.38 ± 0.20 |
| dlPFC [40, 42, 26] | 1.04 ± 0.29 | 2.03 ± 0.35 |

Individual beta estimates were first extracted from ROIs at the time of choice during the dietary decision-making task. Error values correspond to the standard error of the mean. dACC – dorsal anterior cingulate cortex; dlPFC – dorsolateral prefrontal cortex.

SI Table 6. Multilevel General Linear Model of stimulus value for N = 172 participants.

First-level results: Decreased-hunger suggestion group (N = 88)

| | intercept | TR | HR | trial | TRxHR | trialxTR | trialxHR |
|---|---|---|---|---|---|---|---|
| Coefficient | -0.69 | 0.54 | -0.01 | -3.76e-4 | 0.02 | -2.81e-5 | 3.26e-4 |
| STE | 0.06 | 0.03 | 0.02 | 1.71e-4 | 0.01 | 1.23e-4 | 1.21e-4 |
| t | -10.96 | 16.42 | -0.37 | -2.19 | 2.30 | -0.23 | 2.69 |
| Z | -8.21 | 8.21 | -0.37 | -2.15 | 2.26 | -0.23 | 2.62 |
| p | 2.22e-16 | 2.22e-16 | 0.71 | 0.03 | 0.02 | 0.82 | 0.01 |

First-level results: Increased-hunger suggestion group (N = 84)

| | intercept | TR | HR | trial | TRxHR | trialxTR | trialxHR |
|---|---|---|---|---|---|---|---|
| Coefficient | -0.50 | 0.67 | -0.05 | -6.06e-4 | 0.02 | -2.15e-5 | 3.15e-4 |
| STE | 0.06 | 0.03 | 0.02 | 0.00 | 0.01 | 1.37e-4 | 1.34e-4 |
| t | -9.05 | 24.64 | -2.58 | -2.67 | 2.85 | -0.16 | 2.34 |
| Z | -7.53 | 8.21 | -2.52 | -2.61 | 2.78 | -0.16 | 2.29 |
| p | 5.17e-14 | 2.22e-16 | 0.01 | 0.01 | 0.01 | 0.88 | 0.02 |

Second-level, random effects results using weighted two-sample, two-tailed t-tests to compare individual beta estimates from the first level between increased- and decreased-hunger suggestion groups

| | t | df | p | Lower 95% CI | Upper 95% CI |
|---|---|---|---|---|---|
| intercept | -2.28 | 170 | 0.02 | -0.36 | -0.026 |
| TR | -2.81 | 170 | 0.01 | -0.20 | -0.036 |
| HR | 1.08 | 170 | 0.28 | -0.03 | 0.10 |
| trial | 1.39 | 170 | 0.17 | -1.73e-4 | 9.89e-4 |
| TRxHR | -1.02 | 170 | 0.31 | -0.04 | 0.01 |
| trialxTR | -0.82 | 170 | 0.42 | -5.78e-4 | 2.40e-4 |
| trialxHR | 0.85 | 170 | 0.40 | -2.22e-4 | 5.58e-4 |

TR – Taste Ratings; HR – Health Ratings; STE – Standard Error.

SI Table 7. Differential brain activation in response to stimulus value at the time of food choice for the contrast increased > decreased hunger suggestion.

| Region | Brodmann Area | Side | Size | x | y | z | Peak z-score |
|---|---|---|---|---|---|---|---|
| Insula* | 13 | L | 4632 | -30 | -14 | 18 | 4.15 |
| | 13 | R | 271 | 38 | -4 | 18 | 4.51 |
| vmPFC* | 10/11 | | 691 | 0 | 52 | -12 | 4.42 |
| Cerebellum* | | R | 2652 | 12 | -46 | 16 | 4.26 |

| | | | | | | | |
|---|---|---|---|---|---|---|---|
| Precuneus* | 7 | R | 580 | 2 | -42 | 62 | 3.76 |
| NAcc | 25 | | 47 | 0 | 6 | -6 | 3.60 |
| PCC | 31 | L | 115 | -8 | -32 | 52 | 3.45 |

vmPFC – ventromedial prefrontal cortex; NAcc – nucleus accumbens; PCC – posterior cingulate cortex. Values are reported at $p < 0.001$ uncorrected. Regions marked with an * survived $p_{FWE} < 0.05$, family-wise error correction based on the cluster.

SI Table 8. Positive effect of tastiness at the time of food choice for all 57 participants.

| Region | Brodmann Area | Side | Size | x | y | z | Peak z-score |
|---|---|---|---|---|---|---|---|
| IpS | 39 | L | 1122 | -34 | -62 | 38 | 5.89 |
| dACC | 32 | L | 420 | -6 | 34 | 32 | 5.77 |
| dlPFC | 9 | L | 816 | -32 | 20 | 50 | 5.64 |
| | 46 | L | 133 | -44 | 42 | 18 | 5.40 |
| vmPFC | 10 | L | 260 | -6 | 52 | -4 | 5.33 |
| Precuneus | 17/18 | L | 526 | -6 | -64 | 32 | 5.26 |
| PCC | 31 | L | 75 | -4 | -38 | 36 | 4.91 |

IpS – intraparietal sulcus; dACC – dorsal anterior cingulate cortex ; dlPFC – dorsolateral prefrontal cortex; vmPFC – ventromedial prefrontal cortex; PCC – posterior cingulate cortex. Values were taken at $p_{FWE} < 0.05$, family-wise error correction for multiple comparisons based on peak height.

SI Table 9. Negative effect of healthiness at the time of food choice for N = 57 participants.

| Region | Brodmann Area | Side | Size | x | y | z | Peak z-score |
|---|---|---|---|---|---|---|---|
| V1* | 18 | L | 4886 | -16 | -98 | 6 | 6.88 |
| Amygdala* | | L | 308 | -22 | -8 | -18 | 5.31 |
| Parahippo* | 37 | R | 438 | 26 | -48 | -14 | 4.85 |
| vmPFC | 10 | | 2610 | 0 | 64 | 0 | 4.18 |
| STG | 22 | R | 112 | 60 | -10 | -12 | 3.98 |
| vlPFC | 47 | L | 194 | -38 | 28 | -4 | 3.97 |
| PCC | 23 | L | 76 | -16 | -50 | 6 | 3.79 |
| NAcc | 25 | L | 64 | -2 | 14 | -8 | 3.59 |
| MFG | 8 | L | 50 | -16 | 28 | 38 | 3.53 |

V1 – primary visual cortex; Parahippo – parahippocampus; vmPFC – ventromedial prefrontal cortex; STG – superior temporal gyrus; vlPFC – ventrolateral prefrontal cortex; PCC – posterior cingulate cortex; NAcc – nucleus accumbens; MFG – medial forntal gyrus. Values are reported at $p < 0.001$, uncorrected, extent threshold k = 45, regions marked with an * survived family-wise error correction ($p_{FWE} < 0.05$) based on peak height. Note, no significant activation was found for positive correlations with healthiness, even at $p < 0.001$, uncorrected.

SI Table 10. Seeds for parameter estimations

| Inits | $\mu_\alpha$ | $\sigma_\alpha$ | $\mu_{time}$ | $\sigma_{time}$ | $\mu_\theta$ | $\sigma_\theta$ | $\mu_{whealth}$ | $\sigma_{whealth}$ | $\mu_{wtaste}$ | $\sigma_{wtaste}$ | $\mu_{bias}$ | bias. kappa | RNG | RNG. seed |
|---|---|---|---|---|---|---|---|---|---|---|---|---|---|---|
| 1 | 0.5 | 0.05 | 0.5 | 0.05 | 0.1 | 0.05 | 0.3 | 0.05 | 0.01 | 0.05 | 0.4 | 1 | Super-Duper | 99999 |
| 2 | 0.3 | 0.05 | -0.5 | 0.05 | 0.2 | 0.05 | 0.3 | 0.05 | 0.1 | 0.05 | 0.5 | 1 | Wichmann-Hill | 1234 |
| 3 | 0.2 | 0.05 | 0.5 | 0.05 | 0.15 | 0.05 | 0.5 | 0.05 | 0.3 | 0.05 | 0.6 | 1 | Mersenne-Twister | 6666 |

$\mu$ – seed for the hyper-condition parameter mean
$\boldsymbol{\sigma}$ – seed for the hyper-condition parameter precision or error term
bias.kappa – seed weight of the shape and rate of the gamma prior distribution for the participant level bias parameter

SI Table 11. Parameter recovery. Pearson's correlation coefficients between generating and recovered parameters from the tDDM

| Parameters | ALL N=172 |
|---|---|
| $w_{tastecp} - w_{healthcp}$ | 0.79, $p < 2.2e-16$, 95% CI [0.736 – 0.843] |
| $time_{cp}$ | 0.275, $p = 0.0001$, 95%CI [0.137 – 0.403] |
| $bias_{cp}$ | 0.66, $p < 2.2e-16$, 95% CI [0.57 – 0.734] |
| Non-decision time, $\theta_{cp}$ | 0.97, $p < 2.2e-16$, 95% CI [0.96 – 0.978] |
| threshold, $\alpha_{cp}$ | -0.06, $p = 0.40$, 95% CI [-0.203 – 0.08] |
| $w_{tastecp}$ | 0.59, $p < 2.2e-16$, 95% CI [0.4992 – 0.684] |
| $w_{healthcp}$ | 0.67, $p < 2.2e-16$, 95% CI [0.592 – 0.748] |

SI Table 12. Comparison of the posterior distribution means for each population level free parameter of the tDDM model estimated using JAGS and Bayesian approximation of the drift rate

| | **mean($PD_{increased} - PD_{decreased}$)** | **PP** | **95% HDI** | **H1** |
|---|---|---|---|---|
| $\mathbf{w_{healthiness}}$ | 0.17 ± 0.12* | 0.95* | [-0.26 – 0.83]* | Decreased > Increased |
| $\mathbf{w_{tastiness}}$ | 0.19 ± 0.08 | 0.99 | [-0.54 – 0.15] | Decreased < Increased |
| $\boldsymbol{\theta}$ | 0.02 ± 0.04 | 0.31 | [-0.19 – 0.13] | Decreased < Increased |
| $\boldsymbol{\alpha}$ | 0.004 ± 0.07 | 0.50 | [-0.27 – 0.23] | Decreased < Increased |
| **Bias** | 0.04 ± 0.03 | 0.08 | [-0.16 – 0.09] | Decreased < Increased |
| **Time** (seconds) | 0.25 ± 0.23 | 0.86 | [-1.2 – 1.02] | Decreased < Increased |

PD – posterior distributions of each parameter for decreased and increased suggestion groups. PP – posterior probability that differences are < zero (> zero for $w_{healthiness}$). w – drift weights, Ƭ – non-decision time; α – threshold boundary separation, Time – relative starting time.
Note, the * indicates that for the healthiness drift weight the average PDs and the PP were calculated assuming ($d_{decreased} - d_{increased} > 0$).

Table 12a. Summary of the posterior distributions of free parameters from the tDDM obtained using JAGS and a Bayesian approximation of the drift rate

| Decreased-hunger suggestion groups | | | | | |
|---|---|---|---|---|---|
| **Behavioral pilot participants** | | | | | |
| | Lower 95 | median | Upper 95 | mean | SD |
| $w_{healthiness}$ | -0,19 | -0,08 | 0,02 | -0,08 | 0,05 |
| $w_{tastiness}$ | 0,44 | 0,55 | 0,67 | 0,55 | 0,06 |
| $\theta$ | 0,20 | 0,25 | 0,30 | 0,25 | 0,02 |
| α | 0,76 | 0,83 | 0,89 | 0,83 | 0,03 |
| Bias | 0,38 | 0,41 | 0,45 | 0,41 | 0,02 |
| Time | 0,23 | 0,48 | 0,81 | 0,50 | 0,15 |
| **fMRI participants** | | | | | |
| | Lower 95 | median | Upper 95 | mean | SD |
| $w_{healthiness}$ | -0,10 | 0,01 | 0,12 | 0,01 | 0,05 |
| $w_{tastiness}$ | 0,49 | 0,60 | 0,71 | 0,60 | 0,06 |
| $\theta$ | 0,63 | 0,68 | 0,73 | 0,68 | 0,03 |
| α | 0,91 | 0,97 | 1,04 | 0,97 | 0,03 |
| Bias | 0,41 | 0,46 | 0,51 | 0,46 | 0,02 |
| Time | 0,01 | 0,04 | 0,10 | 0,05 | 0,03 |

| Increased-hunger suggestion groups | | | | | |
|---|---|---|---|---|---|
| **Behavioral pilot participants** | | | | | |
| | Lower 95 | median | Upper 95 | mean | SD |
| $w_{healthiness}$ | -0,28 | -0,18 | -0,09 | -0,19 | 0,05 |
| $w_{tastiness}$ | 0,67 | 0,75 | 0,84 | 0,75 | 0,04 |
| $\theta$ | 0,22 | 0,27 | 0,33 | 0,27 | 0,03 |
| α | 0,82 | 0,88 | 0,95 | 0,88 | 0,03 |
| Bias | 0,42 | 0,45 | 0,48 | 0,45 | 0,01 |
| Time | 0,39 | 0,60 | 0,83 | 0,60 | 0,11 |
| **FMRI participants** | | | | | |
| | Lower 95 | median | Upper 95 | mean | SD |
| $w_{healthiness}$ | -0,44 | -0,22 | 0,00 | -0,22 | 0,12 |
| $w_{tastiness}$ | 0,66 | 0,78 | 0,89 | 0,78 | 0,06 |
| $\theta$ | 0,64 | 0,69 | 0,74 | 0,69 | 0,03 |
| α | 0,87 | 0,93 | 0,98 | 0,93 | 0,03 |
| Bias | 0,46 | 0,50 | 0,54 | 0,50 | 0,02 |
| Time | 0,04 | 0,44 | 0,72 | 0,44 | 0,17 |

SI Table 13. Free parameters of the tDDM obtained using deoptim and stepwise approximation of the drift rate

| | **Decreased** | **Increased** | **$BF_{10}$** | **t(170)** | **p** | **Lower CI** | **Upper CI** |
|---|---|---|---|---|---|---|---|
| $w_{healthiness}$ | 0.04 ± 0.08 | -0.08 ± 0.07 | 0.29 | -1.09 | 0.28 | -0.32 | 0.09 |

| | | | | | | | |
|---|---|---|---|---|---|---|---|
| $w_{tastiness}$ | 0.74 ± 0.07 | 1.03 ± 0.05 | 24.73 | 3.32 | 0.001 | 0.12 | 0.46 |
| $\theta$ | 0.66 ± 0.03 | 0.66 ± 0.03 | 0.17 | -0.09 | 0.93 | -0.08 | 0.07 |
| $\alpha$ | 1.73 ± 0.07 | 1.86 ± 0.07 | 0.37 | -1.31 | 0.19 | -0.31 | 0.06 |
| Bias | -0.14 ± 0.03 | -0.08 ± 0.02 | 0.62 | -1.69 | 0.09 | -0.13 | 0.01 |
| Time | 0.21 ± 0.05 | 0.36 ± 0.05 | 2.20 | -2.37 | 0.02 | -0.28 | -0.03 |

The means for the free parameters of the tDDM were calculated for individual parameters in each suggestion group with ± errors corresponding to the standard errors of the mean. Bayesian and frequentist independent sample t-tests compared the two suggestion groups between individual parameters, with a two-tailed null hypothesis (H0: decreased- = increased-hunger suggestion group). $BF_{10}$ – Bayesian independent samples t-test; t(170), p, and 95% confidence intervals (CI) – Frequentist independent samples t-values with degrees of freedom = 170. w – drift weights, $\theta$ – non-decision time, α – threshold boundary separation, Time – relative starting time

SI Table 14. Whole brain activation for the psychophysiological interaction with the vmPFC as a seed ROI (N = 57)

| Region | Brodmann Area | Side | Size | x | y | z | Peak z-score |
|---|---|---|---|---|---|---|---|
| SMG | 40 | L | 843 | -56 | -40 | 46 | 6.20 |
| | 40 | R | 374 | 56 | -36 | 44 | 5.41 |
| Insula | 13 | R | 80 | 34 | 4 | 8 | 5.56 |
| | 13 | L | 199 | -42 | 2 | 4 | 5.26 |
| STG | 22 | R | 66 | 64 | -38 | 16 | 5.28 |
| dlPFC | 46 | L | 73 | -50 | 42 | 8 | 5.02 |
| Premotor/SMA | 44/6 | R | 99 | 54 | 12 | 14 | 4.97 |
| | 6 | R | 52 | -58 | 6 | 16 | 4.97 |
| Fusiform | 37 | R | 65 | 56 | -56 | -8 | 4.90 |
| IpS | 39/7 | R | 45 | 38 | -44 | 42 | 4.86 |
| Putamen | | L | 54 | -24 | -4 | 6 | 4.76 |

SMG – supramarginal gyrus; STG – superior temporal gyrus; dlPFC – dorsolateral prefrontal cortex; premotor – premotor cortex; SMA – supplementary motor area; IpS – intraparietal sulcus.
Values are taken at $p_{FWE} < 0.05$, family-wise error corrected based on peak height.

SI Table 15. Sociodemographic, body composition, and questionnaire data

| | **Decreased hunger** | **Increased hunger** |
|---|---|---|
| Age (years) | 35.67 ± 1.43 | 32.67 ± 1.42 |
| Education | 3.82 ± 0.06 | 3.86 ± 0.04 |
| Weight (kg) | 63.77 ± 1.64 | 63.34 ± 1.38 |
| BMI (kg/m²) | 23.38 ± 0.56 | 23.07 ± 0.44 |
| Body Fat (%) | 28.95 ± 1.11 | 27.93 ± 1.08 |
| SCOFF | 1.37 ± 0.27 | 1.32 ± 0.15 |
| BDI | 4.53 ± 0.056 | 3.88 ± 0.50 |
| YFAS | 0.82 ± 0.22 | 0.86 ± 0.27 |
| IPAQ-sf | 1625.70 ± 181.94 | 1359.92 ± 196.28 |

Note, education was coded as 1 for undergraduate, 2 for high school graduate, 3 for two years of higher education, and 4 for more than two years of university education. SCOFF, YFAS, and IPAQ average scores were calculated for the fMRI study participants only (N = 57). SCOFF – Sick, Control, One stone, Fat, Food; BDI – Beck's Depression Inventory, YFAS – Yale Food Addiction Scale, IPAQ-sf – International Physical Activity Questionnaire-short form
